# *Staphylococcus pasteuri* isolates (*Spvs*) from human atherosclerotic plaques mediate virulence, intracellular resistance and transendothelial invasion of macrophages: A mechanistic paradigm for microbial pathogenicity in atherosclerosis

**DOI:** 10.1101/721571

**Authors:** Nasreen S. Haque, Jayesh Modi, Niloufar Haque, Igor Laskpwski, Romeo B. Mateo, Sateesh C. Babu, Samuel Barasch, John Fallon, Solomon Amar

**Affiliations:** New York Medical College, Valhalla, NY 10595; MidHudson Regional Hospital, 241 North Road, Poughkeepsie, NY 12601; Westchester Medical Center, 100 Woods Rd., Valhalla, NY 10595; Department of Biological Sciences, New York City College of Technology, City University of New York, Brooklyn, NY 11201,USA

## Abstract

Atherosclerotic cardiovascular disease, a chronic inflammatory condition of multifactorial etiology, is the leading cause of death worldwide. The concept of ‘bacterial persistence’ has been proposed as one of several contributing factors to this disease. We hypothesized that the infectious agent(s) found at the site of the atheroma may be dormant but perpetuate virulence in response to host defense and other physiological triggers. In this study, we sought to identify the source of persistent infection in human atherosclerotic plaque and define how pathogen virulence and host defenses mediate plaque vulnerability. Whole genome sequencing (WGS) was used to identify bacteria from pure cultures obtained from atherosclerotic tissues of living subjects diagnosed with more than 70% occlusion of the carotid artery undergoing carotid endarterectomy (CE). WGS identified the predominant species as *S. pasteuri (Spv)* in all CE isolates grown in pure culture except one isolate which was identified as *B. licheniformis*. All *S. pasteuri* isolates (*Spvs*) were found to contain genes for widespread virulence, invasion, and intracellular resistance. As macrophages (Mφs) play a decisive role at all stages of this disease we treated mouse Mφs (RAW 264.7) with *Spvs*. While all *Spvs* tested demonstrated their ability to survive phagocytosis, the highly virulent *Spvs* also activated Mφs and induced trans-endothelial cell migration of these cells which was mediated via the CC chemokine CCL1 and its receptor CCR8. In conclusion, we show that *Spvs* are found in CE plaques, have the capacity to survive phagocytosis and induce the transmigration of Mφs across an endothelial barrier in a CCL1-CCR8 dependent process. These findings highlight the significance of carotid vessels as a reservoir for *S. pasteuri*, a pathogen found in products used routinely in human consumption; this may explain how microbial pathogenicity modulates plaque vulnerability in atherosclerosis.

## Introduction

Atherosclerosis is the number one cause of morbidity and mortality worldwide. This high incidence of atherosclerosis is not fully explained by conventional risk factors (1). While numerous hypotheses have been proposed for this chronic condition, it is widely accepted to be driven by an inflammatory process. One of the most interesting developments in recent years has been the "persistent infection hypothesis" which claims that microbial agent(s) may play an important role in atherogenesis; either through a direct pro-inflammatory effect on the vessel wall or through a less specific, long-distance pro-inflammatory effect (2, 3) The validity of the hypothesis that infection contributes to atherosclerosis has not been firmly established, although the evidence is becoming compelling. Both viral and bacterial populations have been shown to be present in individuals with atherosclerosis (4, 5). While symptomatic infected carotid artery was concurrent with bacterial endocarditis (6) the association of infections such as periodontitis with atherosclerotic diseases is well documented (7, 8 and 9). The development of atherosclerosis is also known to be influenced by certain gut microbes which include species like *Escherichia coli*, *Enterobacter aerogenes*, and members of *Klebsiella spp.* genus. Interestingly, bacterial species such as *Roseburia intestinalis* and *Faecalibacterium prausnitzii*, associated with the production of short chain fatty acids are found to be almost depleted in atherosclerosis patients (10). Oral bacteria such as *Lactobacillus salivarius*, *Solobacterium moorei*, *Atopobium parvulum*, and species belonging to *Streptococcus spp.* genus have been shown to be positively correlated with atherosclerosis when found in the gut (7). In this regard, commensal as well as pathogenic gut microbiome modulates atherosclerotic cardiovascular disease (11, 12 and 13). Recently, trimethylamine-N-Oxide (TMAO) synthesis and bile acid metabolism mediated re-modelling of the gut microbiota has been suggested to play an important player in atherosclerosis by regulating TMAO (14, 15). However, even as several microbial species have been directly found in the atherosclerotic plaque, the association remains inconclusive.

At present, no pathogen(s) have been identified, nor cultured from the atherosclerotic plaque that can inform to the etiology of this disease. We hypothesized that the infectious agent(s) found at in the atherosclerotic niche may be dormant but perpetuate virulence in response to specific environmental triggers. Thus, the identification of infectious agent(s) linking the ‘persister’ phenotype with the virulence induced characteristics may explain the nuances of atherosclerotic plaque initiation, development and/or progression. In the present study, we examined the microbiome of atherosclerotic tissues of living subjects diagnosed with more than 70% occlusion of the carotid artery undergoing carotid endarterectomy (CE) by 16S and whole genome shotgun metagenomic sequencing. Shotgun sequencing of microorganisms has been successfully used to identify the microorganisms both from direct as well as cultivated patient specimen, by both 16S and whole genome sequencing (WGS) and *in silico* analysis of their taxonomy, phylogenetic relationships and potential virulence potential (16, 17). The present study shows that 16S sequencing together with WGS based analysis is suitable for the identification of the potential infectious agents from human CE specimens.

We have identified *S.pasteuri (Spv)* as the dominant species present in human atherosclerotic plaque which contains genes for virulence and resistance and has the ability to mediate intracellular resistance and transendothelial invasion of mouse macrophages (Mφs). In addition, *Spvs* are found to be enriched for genes required for iron sequestration. This is important at two levels. First iron activates oxidation of low density lipoprotein (LDL) which is known to induce endothelial cell injury and recruitment of Mφs (with infiltrated *Spvs*) to the vessel wall (VW). Thus, *Spvs* may initially be tolerated due to their ability to sequester iron/heme, thereby reducing iron overload and limiting the formation of oxidized LDL and inhibiting vascular injury. However, the ability for transendothelial invasion of Mφs places infiltrated *Spvs* at the core of the plaque which upon stressful triggers such as Mφ apoptosis will cause bacterial dissemination leading to colonization and formation of biofilms or bacterial death with release of iron/heme in the environment perpetuating oxidation of LDL and subsequent destabilization of the plaque. These findings highlight the significance of carotid vessels as a reservoir for *S. pasteuri*, a species of concern due to its conserved virulence, ability to persist for long periods (e.g. catheters, biofilms) and presence in products used in human consumption (e.g. fish, milk, sausage, drinking water); which may explain the role of microbial pathogenicity in atherosclerosis. Thus, mapping the trajectories of *S. pasteuri* virulence and host response may predict the fate of a vulnerable plaque and assist in attenuating the progression of this disease.

## Methods

### Isolation of and from human subjects undergoing carotid endarterectomy

Atherosclerotic tissues of living subjects diagnosed with more than 70% occlusion of the carotid artery undergoing carotid endarterectomy (CE) after informed consent had been obtained at the Department of Vascular Surgery, Westchester Medical center, Valhalla, NY. A total of 44 samples from CE from 11 individual patients were examined in this study. Specimen isolation, culture and workflow for sequencing and data analysis are shown (Fig 1, 2). The samples were obtained from April 2016 to August 2018 and further processed under sterile conditions. CE samples from each CE plaque were sub-divided into 2 sections; containing the plaque (n=11) or without plaque as control (n = 7). Each tissue with the plaque was further divided into 4 sections (2 mm each); 2 sections were used to isolate DNA for direct metagenomic sequencing and 2 sections were used to isolate bacteria and grow in pure culture for whole genome sequencing and the remaining material was stored at −80 °C. Similarly for the control sample 2mm sections were used for 16S and WGS shotgun metagenomics analysis.

**Fig 1.**
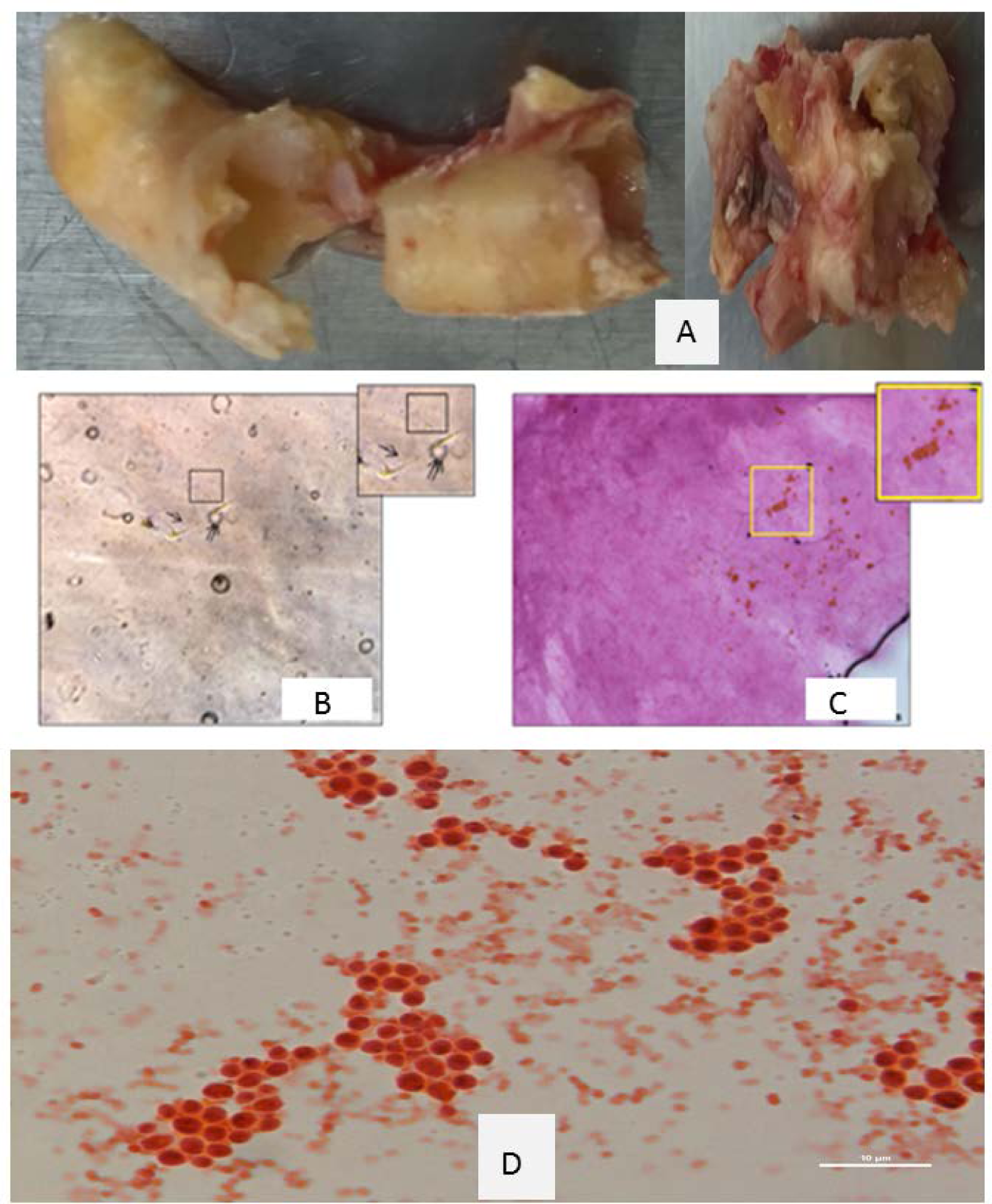
Bacteria from CE plaque were isolated and grown in pure culture. CE plaque extracted tissue is shown (Fig 1A). Bacterial microcolonies were isolated and observed under phase contrast (1B; mag: 20x; inset, 40x) or stained with safranin stain (1C; mag: 20x; inset, 40x). In addition, Gram stain of CE derived colonies grown in pure culture is shown [Fig 1D; mag; 10x].

**Fig 2.**
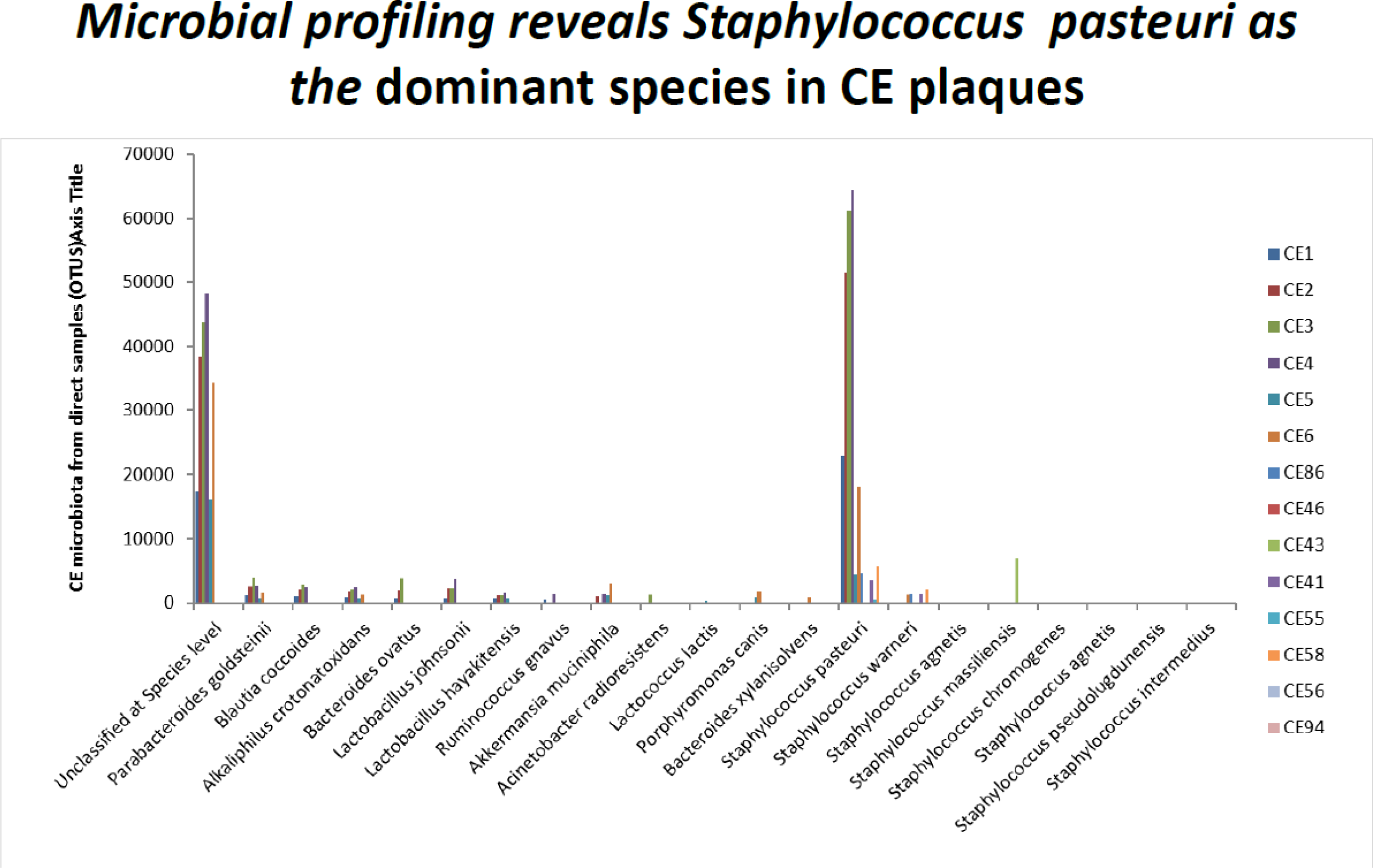
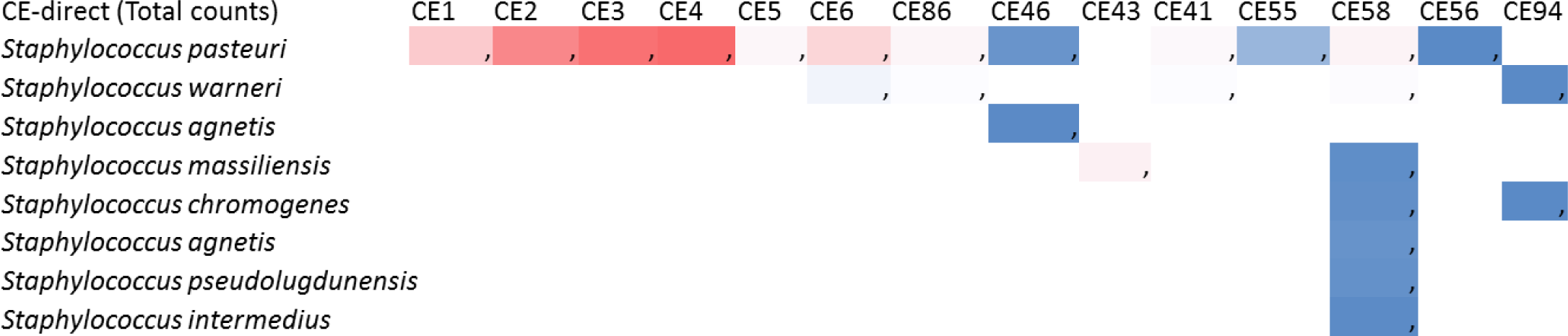
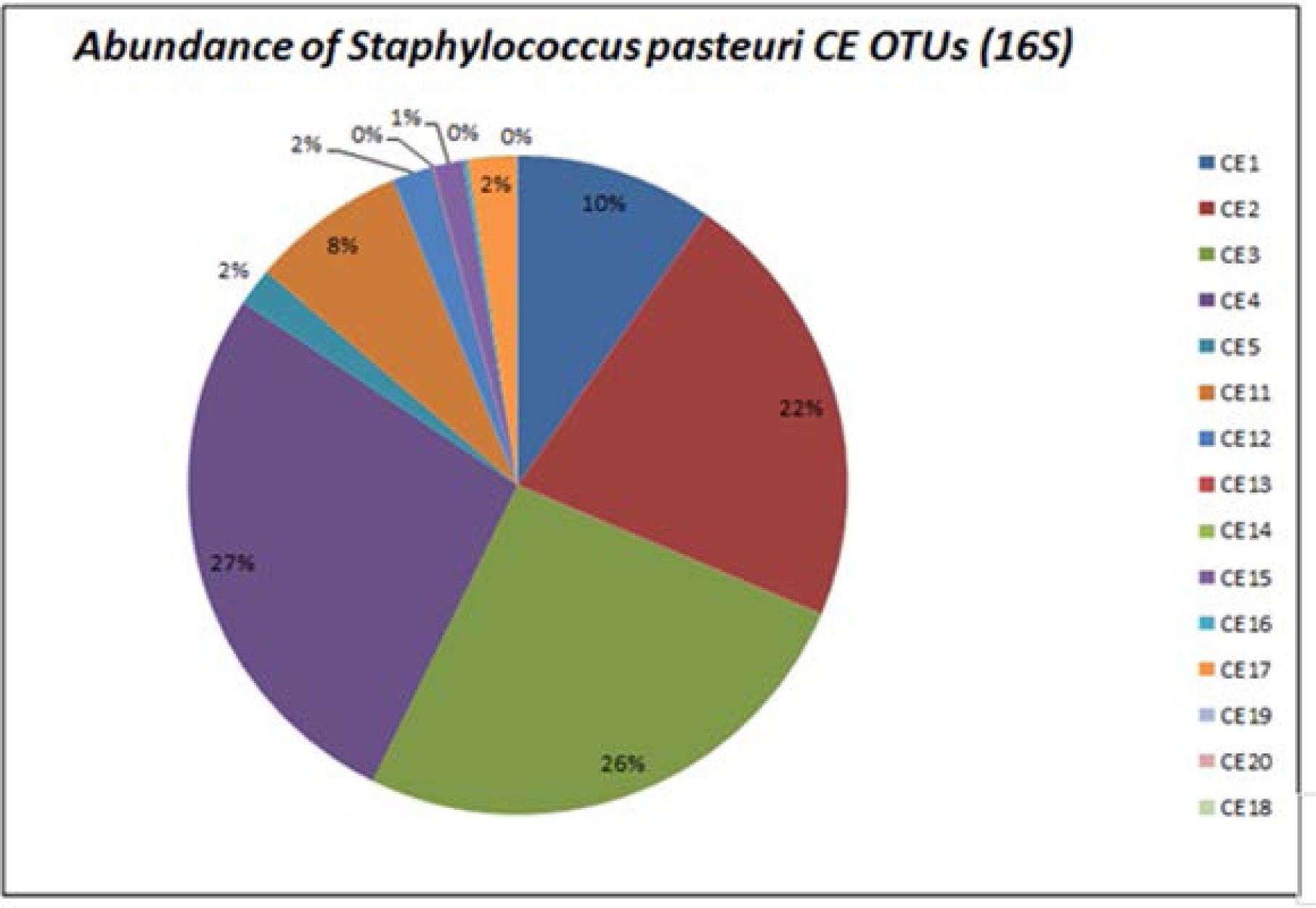
16S metagenomic and whole genome sequencing analysis identifies *Staphylococcus pasteuri in all CE* plaque. Distribution of microbiome in CE plaque identified by direct 16S sequencing is shown at the species level which demonstrated that while each sample contained a unique microbiome, *Staphylococcus sps* (OTUs) dominated in all samples tested (Fig 2A, 2B). WGS further identified the predominant species as *S. pasteuri* in all CE isolates grown in pure culture (Fig 2C) except in one isolate which was identified as *Bacillus licheniformis*.

### Bacterial Cultivation

Aliquots from the vitreous specimens were cultivated for 12 days on 5% horse blood agar, chocolate agar, brain heart infusion broth under anaerobic conditions and on anaerobic plates (SSI Diagnostica, Denmark) under anaerobic conditions at 35 °C were cultivated for 12 days on 5% horse blood agar, chocolate agar, brain heart infusion broth under anaerobic conditions and on anaerobic plates (SSI Diagnostica, Denmark) under anaerobic conditions at 35 °C according to the standard operating procedure. Brain Heart Infusion (BHI) Agar derives it nutrients from the brain heart infusion, peptone and glucose components (BD). To isolate bacteria from CE samples 100 μl aliquots were distributed on Blood Agar (BA) and macconkey agar (MA) (Fischer Scientific). The BA and MA plates were incubated for 3 days at 37 °C under both aerobic and anerobic conditions, colonies were harvested harvested, grown in pure culture and stored in DMSO solution 7% dimethylsulfoxide (vol/vol) at −80 °C. One representative colony morphotype per sample was selected for whole genome sequencing.

### Isolation of DNA and shotgun Metagenomic sequencing

The DNA was prepared and sequenced according to the Nextera XT DNA Library Preparation Guide. DNA was isolated from direct CE specimens as well as from CE derived bacteria growing in pure culture. For each round of DNA isolation, one extraction control (blank) was included. DNA was extracted using QIAGEN’s DNeasy Blood & Tissue kit and after samples have been quantified, sequence libraries were generated following Illumina’s TruSeq DNA preparation protocols. The generated library was further quantified, pooled and sequenced using anIllumina NextSeq550 using paired-end sequencing in our Genomics Core facility, New York Medical College. A total read pairs obtained from the samples were of 18,128,142 ranging between 6,802,761–2,431,211 for the samples from patientsundergoing carotid endarterectomy (CE). Some reads were too short and were removed and not shown. *For whole genome sequence analysis i*solates were sequenced (2 × 150 bp paired-end) on a MiSeq system (Illumina, San Diego, CA, USA) as previously described (18). Reads were adapter trimmed and filtered for phiX reads using BBduk. The high-quality reads where assembled using the SPAdes assembler and the genome sequence assemblies analysed using the bacterial Analysis Pipeline (19, 20).

### Bioinformatics and Data Analysis

The metagenomics analysis was carried out first by adapter and quality trimming and low complexity filtering of raw reads using BBDuk of BBMap version 35.82 (http://jgi.doe.gov/data-and-tools/bbtools/). As atherosclerotic plaques tissue were expected to contain large amounts of human genome DNA which can reduce the microbial genome DNA reads during sequencing the human-affiliated reads was removed from the samples by mapping the reads against the reference genome GRCh38.p10 (GCF_000001405.36) and reads that aligned to human sequences in the non-redundant nucleotide collection (nt) database from NCBI. Detection of ambiguous sequences in public reference genomes and creation of curated microbial genome database.Reads were classified in samples using Kraken followed by Bayesian re-estimation of abundance (Bracken). Classification of reads were performed using BLASTn of BLAST version 2.6.044. For details, see: (Supplementary data: *Spvs* Genome neighbors; Contigs, AR, Virulence, phage). Multiple genome alignments provide a basis for research into comparative genomics and the study of genome-wide evolutionary dynamics. Mauve, (http://darlinglab.org/mauve/mauve.html) is a system for constructing multiple genome alignments in the presence of large-scale evolutionary events such as rearrangement and inversion. We Mauve to align and compare sequence distance and homology between the different contigs obtained from sequencing of CE derived isolates. The data was further processed and after annotation, multiple metagenomic sequence comparisons were made and a functional analysis performed. To obtain a tentative pathway analysis, an analysis based on RAST (Rapid Annotation using Subsystem Technology- http://rast.nmpdr.org/) was performed to investigate metabolic or disease pathways. RAST is a fully-automated service for annotating complete or nearly complete bacterial and archaeal genomes and provides high quality genome annotations for these genomes across the whole phylogenetic tree.

### Ethics

This study was performed in accordance with the Declaration of Helsinki. The study was approved by the US Data Protection Agency and by the local ethics committee of New York Medical College, Valhalla, NY (IRB# L-10,694).

### Data accessibility

The sequencing data generated and analyzed in this study are available from DDBJ/ENA/GenBank. (Note: Will upload this upon the submission of this manuscript).

## Results

### *Staphylococcus pasteuri* dominate in CE plaques

Distribution of microbiome in CE plaque was identified by direct 16S sequencing which is shown at the species level which demonstrated that while each sample contained a unique microbiome, *Staphylococcus sps (*OTUs) dominated in all samples tested (Fig 2A, 2B). The dominance by firmicutes was confirmed in our studies. The microbial population identified by 16 S metagenomic sequencing directly from the CE samples were classified (Fig 2A) as *Parabacteroides goldsteinii, Blautia coccoides, Alkaliphilus crotonatoxidans, Bacteroides ovatus, Ruminococcus gnavus, Akkermansia muciniphila Acinetobacter radioresistens, Lactococcus lactis, Porphyromonas canis, Bacteroides xylanisolvens, Lactobacillus johnsonii and Lactobacillus hayakitensis and Staphylococcus sps. (S. pasteuri, S. warneri, S. agnetis, S. massiliensis, S. chromogenes, S. agnetis, S.pseudolugdunensis S. intermedius)* with a predominance *S. pasteuri* in all CE samples tested (Fig 2B,2C).

### *Staphylococcus pasteuri isolated from from CE* plaque grown in pure culture

We have successfully isolated bacterial colonies from CE plaque from multiple samples and under specific conditions (see Materials and Methods) obtained from atherosclerotic tissues of living subjects diagnosed with more than 70% occlusion of the carotid artery undergoing CE (Fig 1). This suggests that these bacteria may be dormant but can be activated as seen in our *in vitro* studies. A total of 17 colonies were grown in pure culture (clones 1,2,3,4,11,13,14,15,16,17,18,19,20, CE1, CE2, CE3 and CE4). 15 isolates from pure cultures were subjected to WGS. All CE isolates grown in pure culture were identified as having highest sequence homology to *S. pasteuri* except one isolate which was identified as *Bacillus licheniformis (Fig 3B)*. In total 11 isolates [*S. pasteuri variants* (*Spvs*) =10 isolates and 1 *B*. *licheniformis)* =1 isolate] were identified. The closest neighbors to *S. pasteuri*SP1 is shown (Fig S1, S2).

**Fig 3.**
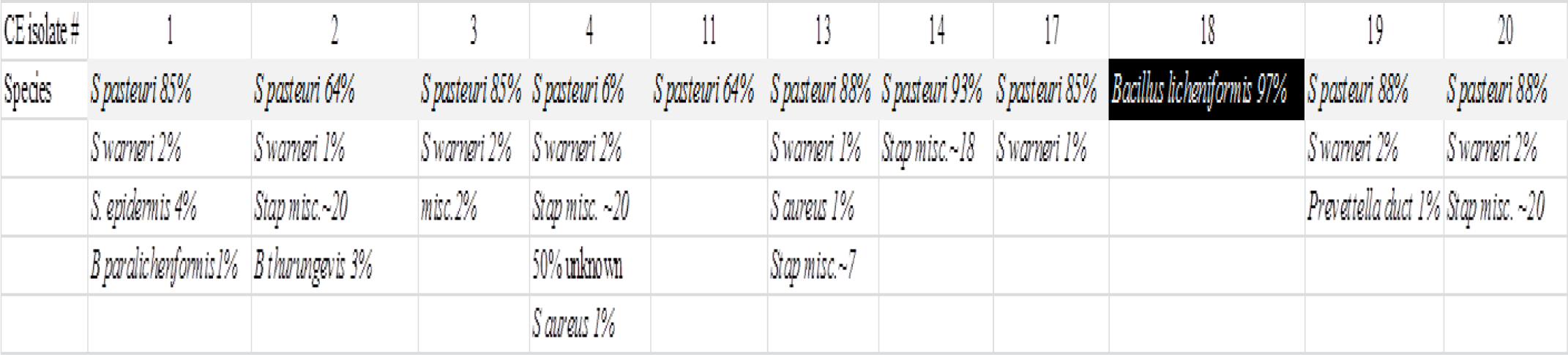
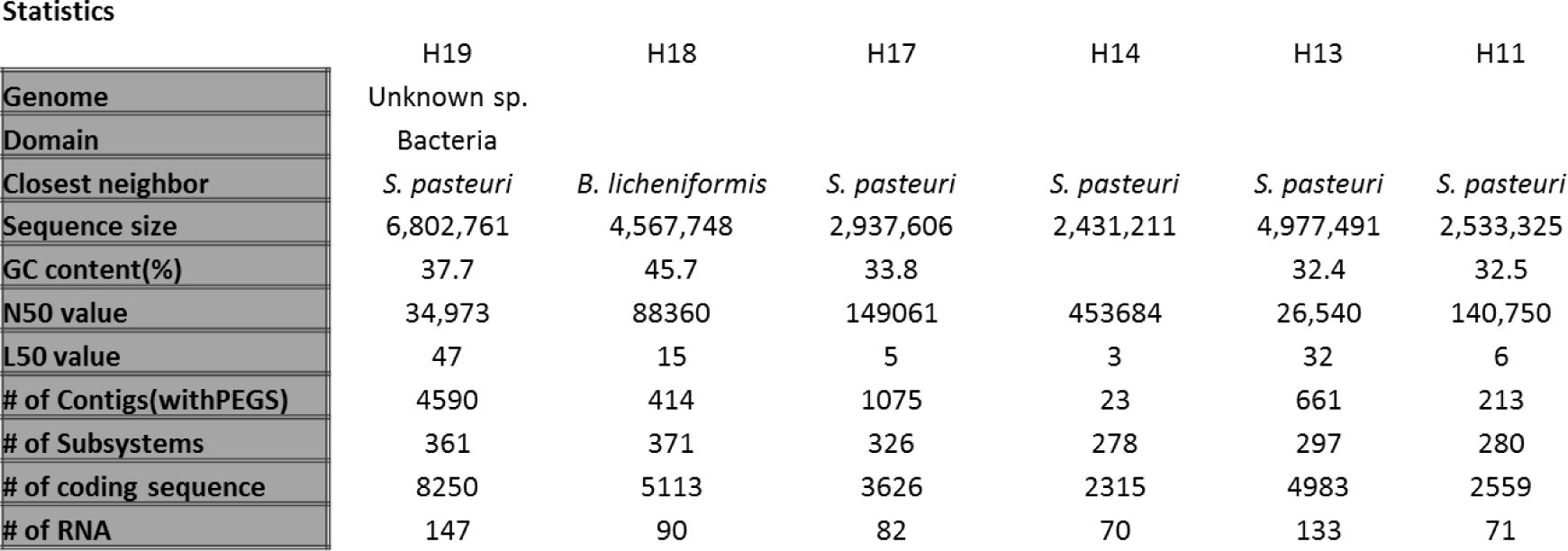
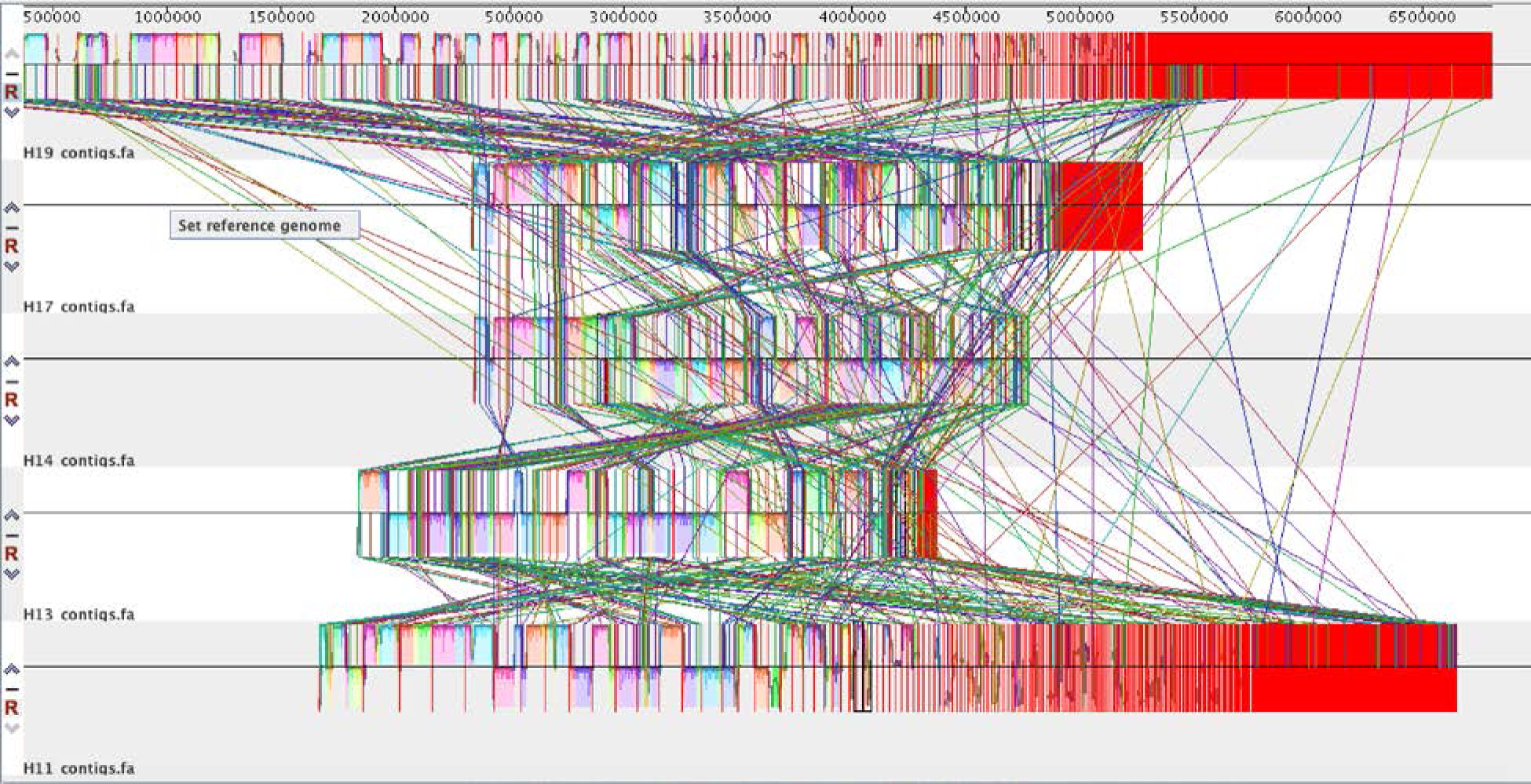
Genomic organization of CE plaque derived isolates (Spvs) Genomic organization of CE plaque isolates (*S. pasteuri* H19, H17, H14, H13, H12 and *B. licheniformis;* H18) was ascertained using online servers RAST (). It was found that H19 had the largest sequence size compared to other Spvs and B. *licheniformis* (Fig 3A, 3B). Genomic size was found to be positively correlated to the coding sequence, number of RNA and subsystems found in each isolate and comparative homology was done by Mauve (3C) *refer to Supplemental data in the results section.

### Genome size is a determinant of frequency of conserved domains in *Spvs*

Genomic organization of CE plaque isolates (*S. pasteuri* H19, H17, H14, H13, H12 and *B. licheniformis;* H18) was further compared using online servers RAST and Mauve. It was found that H19 had the largest sequence size compared to other *Spvs* and B. *licheniformis* (Fig 3B) which was found to be positively correlated to the coding sequence, number of RNA and subsystems found in each isolate. Comparative homology was done by Mauve (3C) which shows the multiple sequencing alignments of the different isolates (Fig 3C). *Spv19* with the largest sequence contained the highest number of functional domains in the subsystem features suggesting that genome size is a determinant of conserved domains in Spvs (Fig 4A). Distribution of genes related to sub functions is shown (Fig 4A, details supplementary data S3-S7). Within *Spv19* the number of genes for cell wall synthesis and phage is highest compared to genes for secondary metabolites suggesting that higher genomic content probably provides for the essential nutrients for maintenance and therefore, secondary metabolites are not essential for their growth, development and reproduction. Interestingly, *Spv13* with comparatively shorter sequence length contained higher number of genes for secondary metabolites (Fig 4B).

**Fig 4.**
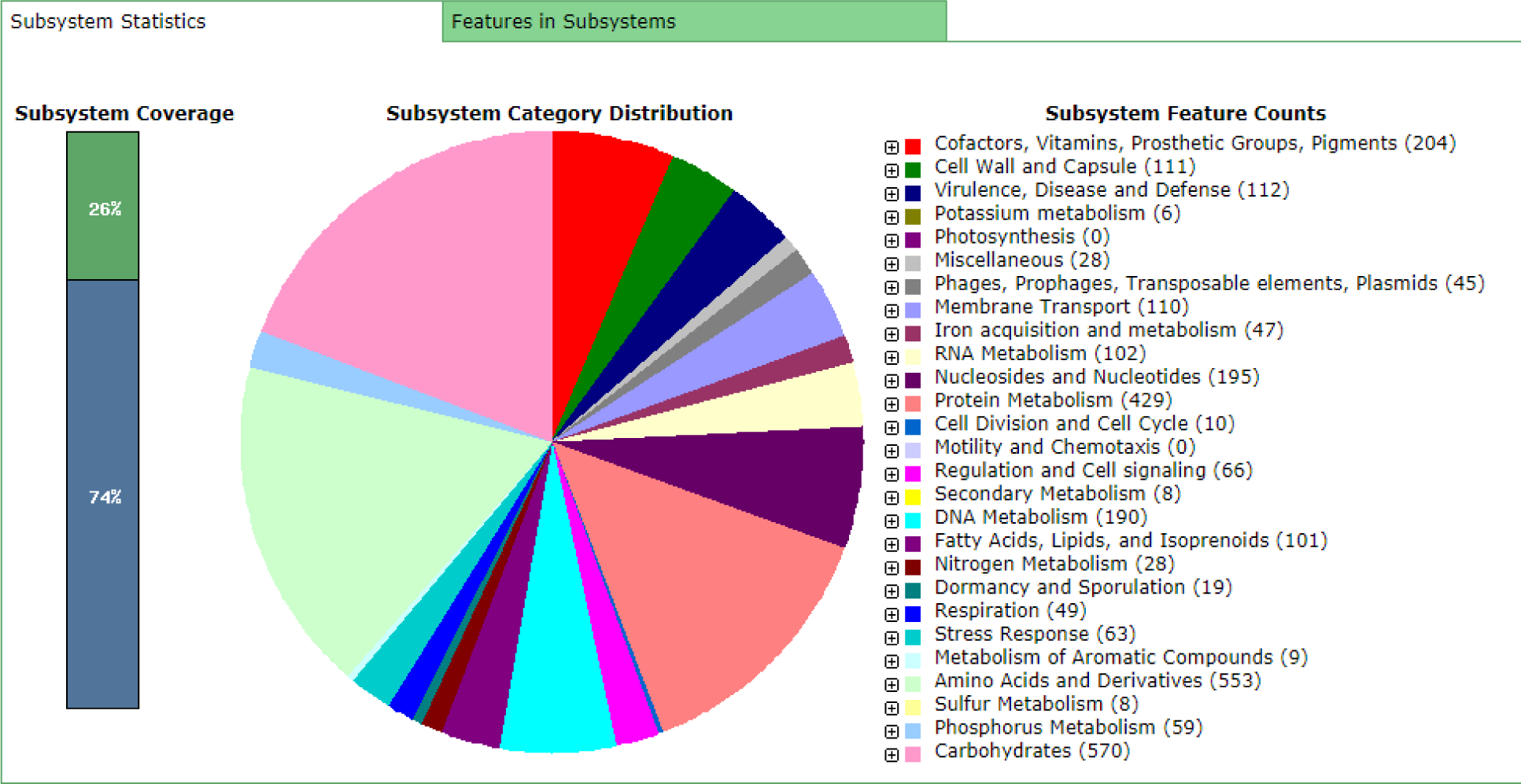
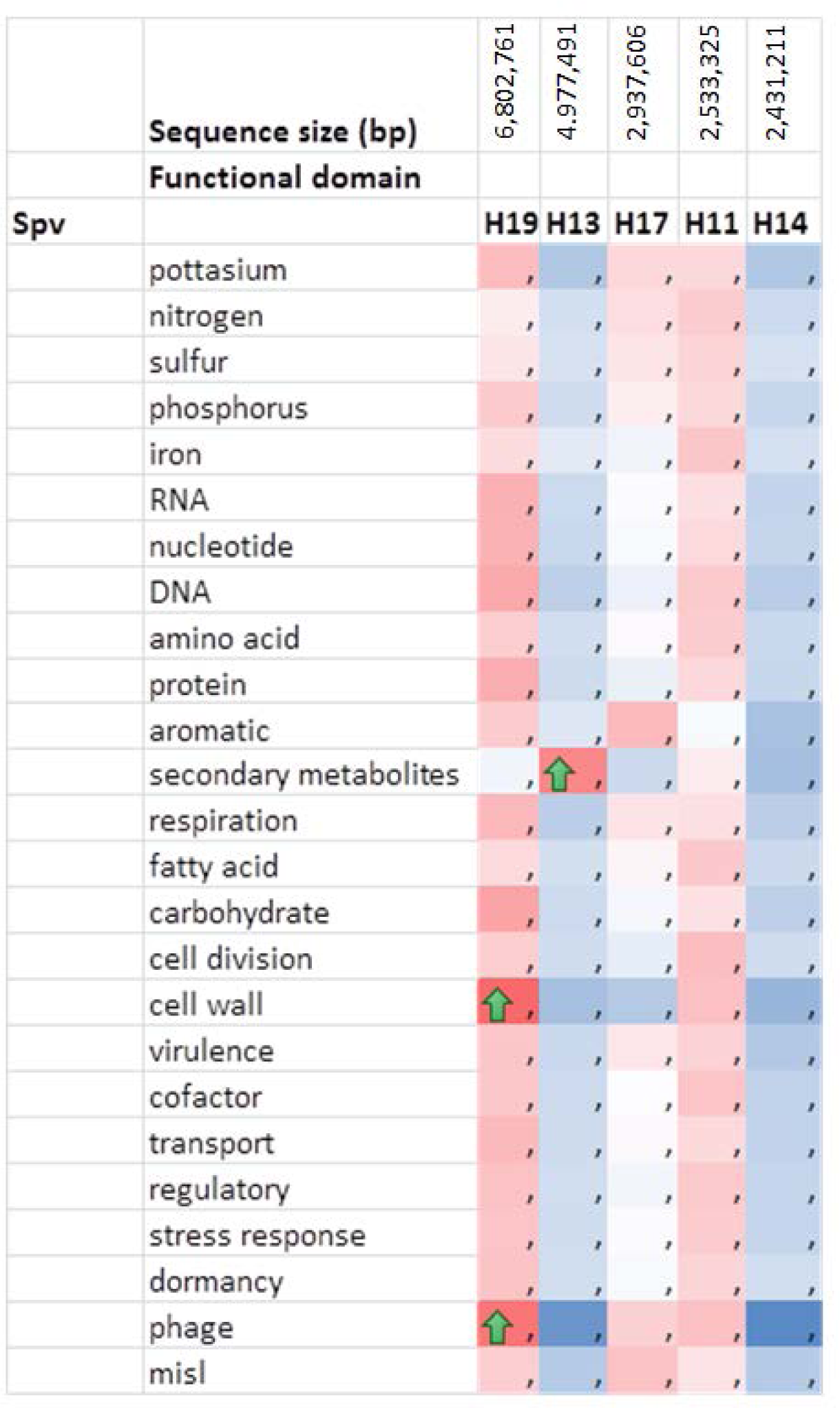
Genome size is a determinant of conserved domains in Spvs. *Spv19* with the largest sequence contained the highest number of functional domains in the subsystem features suggesting that genome size is a determinant of conserved domains in Spvs (Fig 4A,). Within *Spv19* the highest percentage was found in cell wall synthesis and the phage content. Interestingly, there was low number of genes for secondary metabolites which was found to be significantly high for *Spv11* (Fig 4B)

### CE isolates contain genes for virulence, intracellular invasion, resistance and iron acquisition

As Mφs infiltration to the site of injury is the first step towards the development of an atherosclerotic plaque we studied how *Spvs* may modulate this process. Rast database was used for initial analysis of bacterial genome. First, we found that all CE isolates (*Spvs* and *B. licheniformis*) contained genes for virulence, intracellular invasion and resistance (e.g toxic compound, antibiotics, and bacteriophages) and iron acquisition (. In addition, all isolates also possessed genes for bile hydrolysis which is known to also to assist in microbial uptake of iron (Fig 5A). This is important because oxidation of LDL by iron/heme is known to play a key role in the initiation and progression of atherosclerosis. Iron acquisition genes found in Spvs also included siderophore anthrrachelin. Interestingly, we found that genes for iron acquisition were abundant in all isolates (e.g. iron and heme/hemin uptake) were derived from *S. aureus* (5B). Virulence operon acquired invasion and intracellular resistance were derived from *Mycobacterium tuberculosis* virulence operon which included genes for DNA transcription, quinolinate biosynthesis and SSU ribosomal proteins and LSU ribosomal proteins required for protein synthesis (Fig 5C).

**Fig 5.**
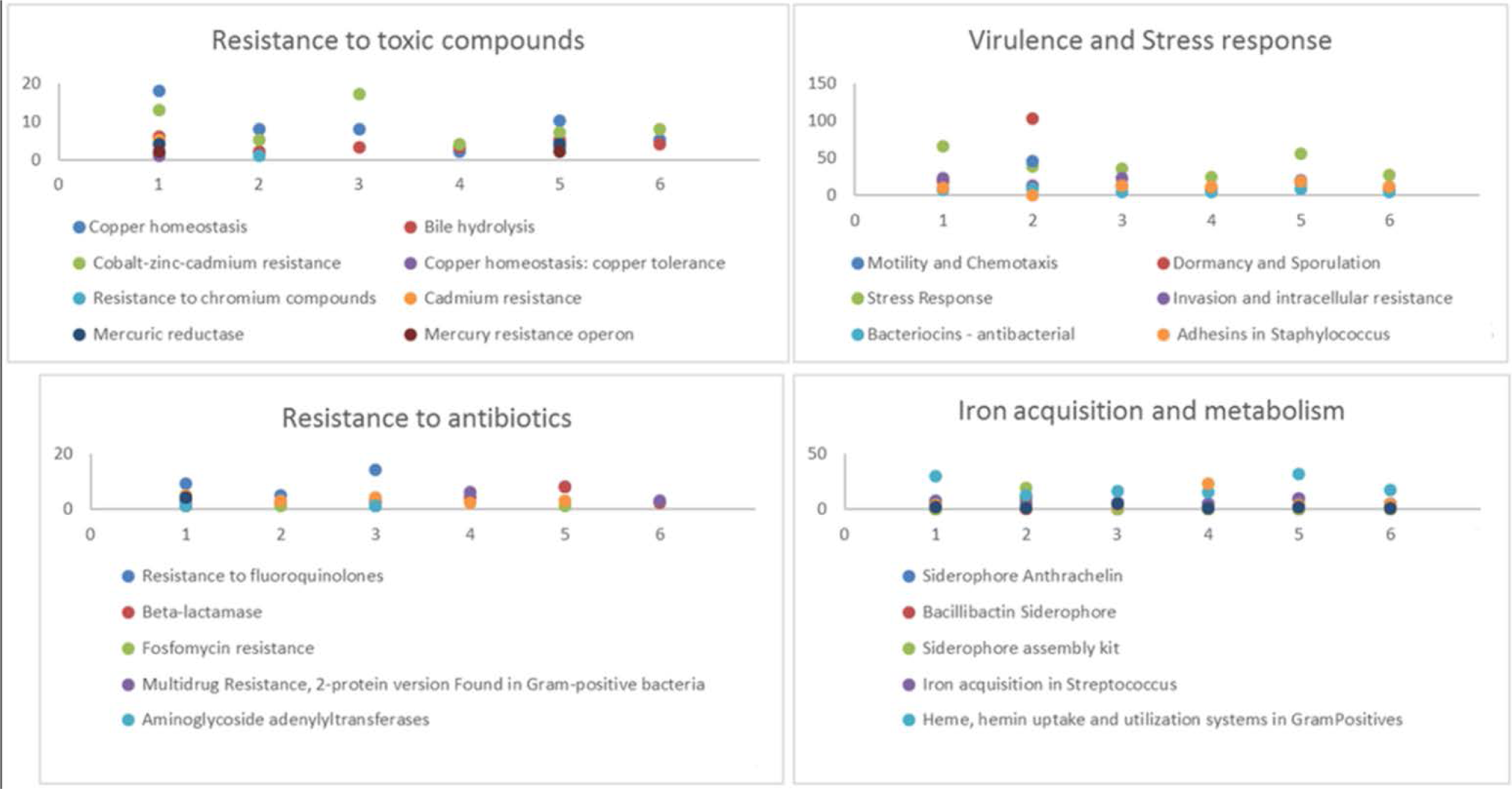
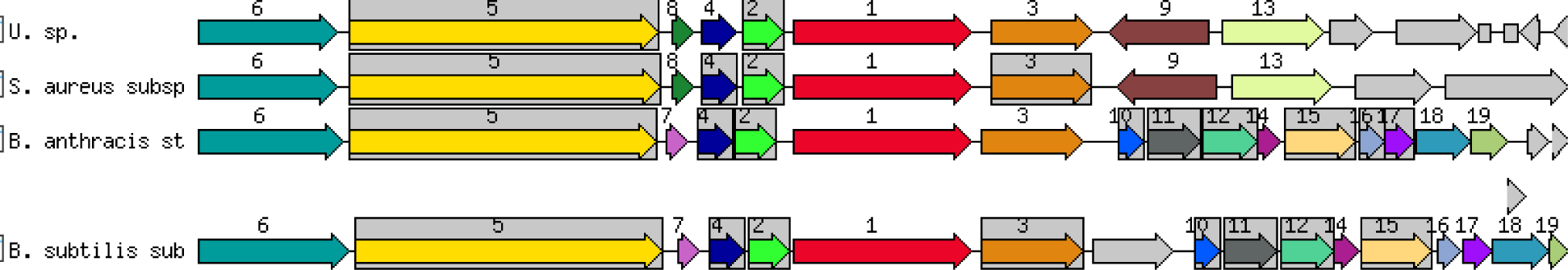
Virulence operon acquired invasion and intracellular resistance and iron acquisition in Spvs. Distribution of genes related to sub functions shown includes resistance to toxic compounds, virulence and stress response, resistance to antibiotics and iron acquisition and metabolism (Fig 5A). Overall, virulence operon acquired invasion and intracellular resistance and genes iron acquisition were abundant in all isolates (e.g. iron and heme/hemin uptake) which is specifically derived from *S. aureus.* Iron acquisition genes found in Spvs also included siderophore anthrrachelin. In addition genes for DNA transcription, quinolinate biosynthesis and protein synthesis (SSU and LSU ribosomal proteins) were found to be acquired from mycobacterium virulence operon (Fig 5B). Number 1 (red) 1 denotes siderophore Anthrrachelin acquired from mycobacterium virulence operon (Fig 5B). In addition DNA transcription (#2, Quinolinate biosynthesis (#5) and protein synthesis(#2,4 [SSU and LSU ribosomal proteins (Fig 5C).

For a more complete analysis of virulence in *Spvs* the data obtained from WGS was further analyzed by using the VRDB database. It was further confirmed that *Spv1* and *Spv19* were highly virulent (Supplemental Fig 2) compared to *Spv11, Spv13* and *Spv17.* While all isolates were found to have virulent genes for (i) growth under stressful conditions(capsule), (ii) phagocytosis resistance(clumping factor), (iii) platelet aggregation, (iv) autolytic activity and proteolytic maturation of SspB, (v) degradation of bacterial cell surface fibronectin binding protein(V8 protease); *Spv17* were found to have the ability to (i) detach and facilitate the spread of infection and cleave human plasma in vitro (Aureolysin), (ii) form biofilm (EbP pilli), detoxify H_2_O_2_ and protects against reactive oxygen species (KatA). KatA was also found in *Spv11*. The following virulent genes were only found in *Spv1 and Spv19*: A 8 gene cluster (TypeVII) which when absenct in staphylococcus reduces virulence, TypeII collagen (Acm), increases cell adhesion(AS), Biofilm production (BopD), resists toxicity of bile and bile salts (BSH), binds collage type V (EcbA), induces Quorum Sensing and regulates bopD, gelE, sprE expression important for biofilm formation and metabolic pathways (Fsr), novel invasion favoring entry into non-professional phagocytes but not in invasion of phagocytes (LpeA), adhesins play an important role in tissue tropism of pathogenic bacteria (Myf/pH6 antigen), mangnesium and zinc; mutants have impact on colony formation and oxidative damage (PsaA), binds Collagen TV and fibronogen (Scm), biofilm medical devise; binds fibronogen and nidogen (SgrA), modulates host immune response; persistance colonization of the trachea (TTSS). In addition, *Spv1* and *Spv19* have acquired a pathogenicity island in Locus of enterocyte effacement LEE (EPEC) which mediates colonization, persistence; attachment and biofilm formation on abiotic surfaces (Esp). enterogenic isolate EPEC is identified in pathogenecity island HZT in highly pathogenic isolate *Spv19. S. pasteuri* isolates in CE plaques are virulent with abundance of *E. coli* derived Pathogenicity Island (Supplemental data S8).

### Antibiotic resistance (AR) is prevalent in all *Spvs*

CE isolates H1, H2, H3, H4 and H11 did not demonstrate AR while all other isolates contained AR to some degree, albeit in varying degrees. Multiple antibiotic resistant were found in all isolates. Percentage coverage for resistance in all *Spvs* was found to be highest for aminoglycoside [(AGly)-APH-Stph], ATP binding efflux pumps (sav1866,arlR), MDR flouroquinine (norA, mgrA) and tetracyclin (Tet) TetU, mepA and trimethropin (dfrE, dfrC, dfrR,dfrF)(Fig 5A, 6A). Interestingly, isolate CE19 which we found to be the most virulent also contained all features required for vancomycin resistance (Fig 6A). CE isolates were further subjected to various antibiotics that are routinely used in the clinic (e.g. penicillin, tetracycline, ampicillin, streptomycin, gentamicin) and the zone of inhibition was measured (Fig 6B). It was found that all isolates showed some degree of susceptibility as well as resistance to different antibiotics tested (Fig 6B). All Spv isolates used in this study showed resistance to gentamicin, as evidenced by the zone of inhibition, except isolate # H18 which was identified as *B. licheniformis*. Interestingly, gentamicin is an aminoglycosides which has been reported in clinical isolates of *Staphylococcus aureus* (21) and in *S pasteuri* (22) suggesting that *Spvs* found at the CE plaque may also be clinically relevant. At least 3 isolates were completely resistant to tetracycline and 1 isolate was resistant to streptomycin while susceptibility to ampicillin, penicillin G, streptomycin and tetracycline was observed at varying range (Fig 6B). Bacteriophage diversity in *Spvs* was also ascertained which showed that all isolates tested contained at least one bacteriophage; however, variation in the type of genes was observed (Fig 6C). The phages belong to the family siphoviridae and in subclass phietalikevirus with representatives from staphylococcus, streptococcus and proteus genera. In addition, two isolates showed the presence of fungal gene.

**Fig 6.**
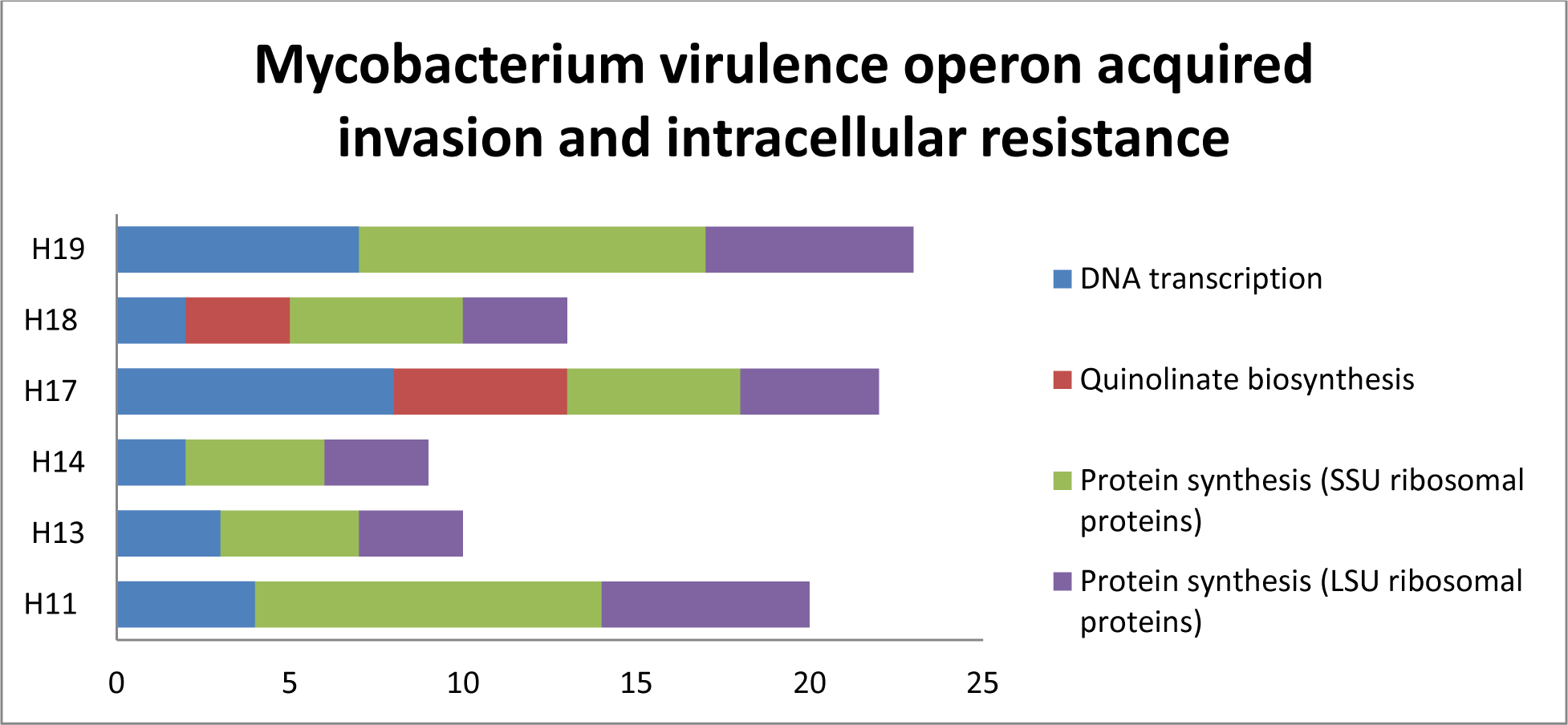
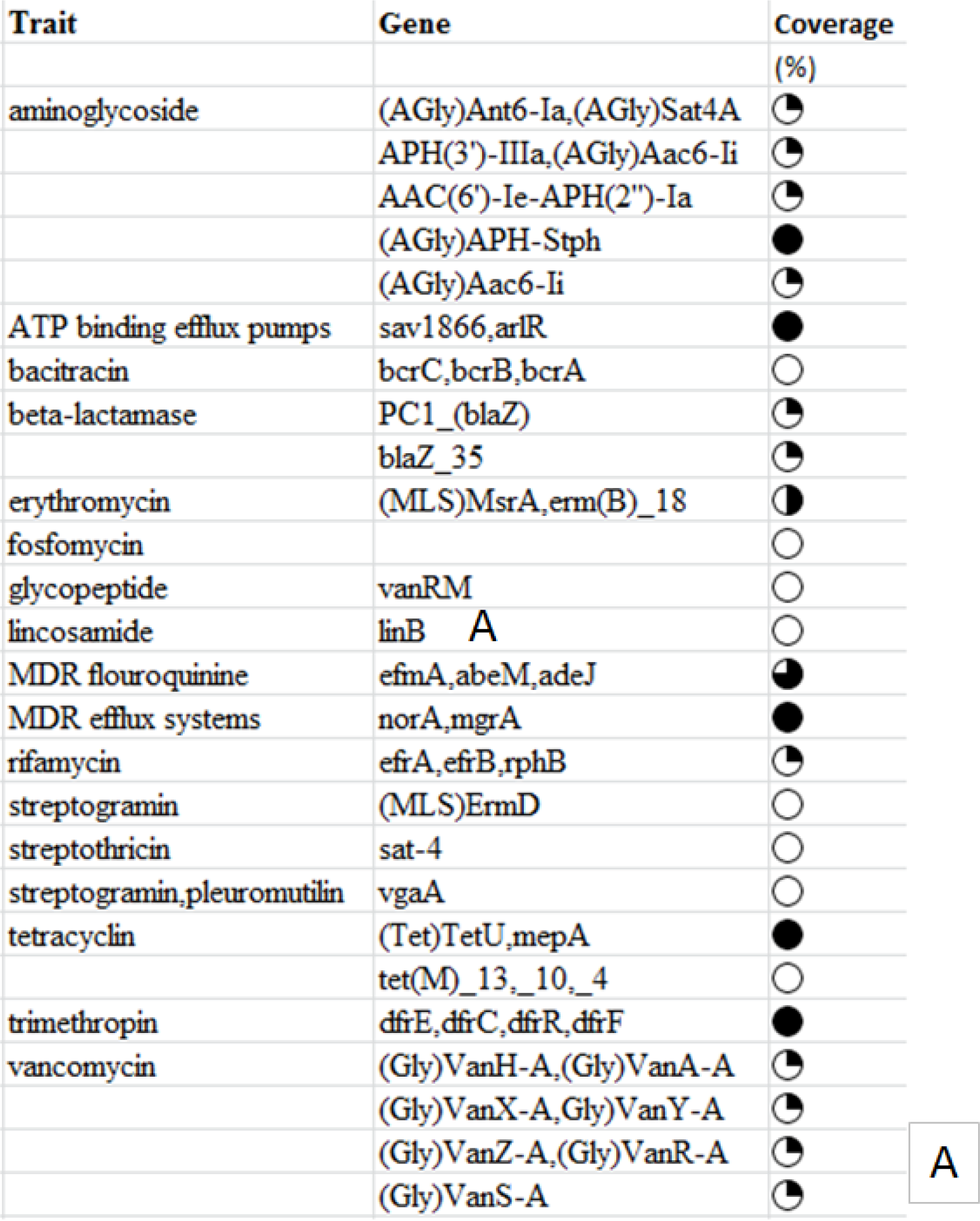
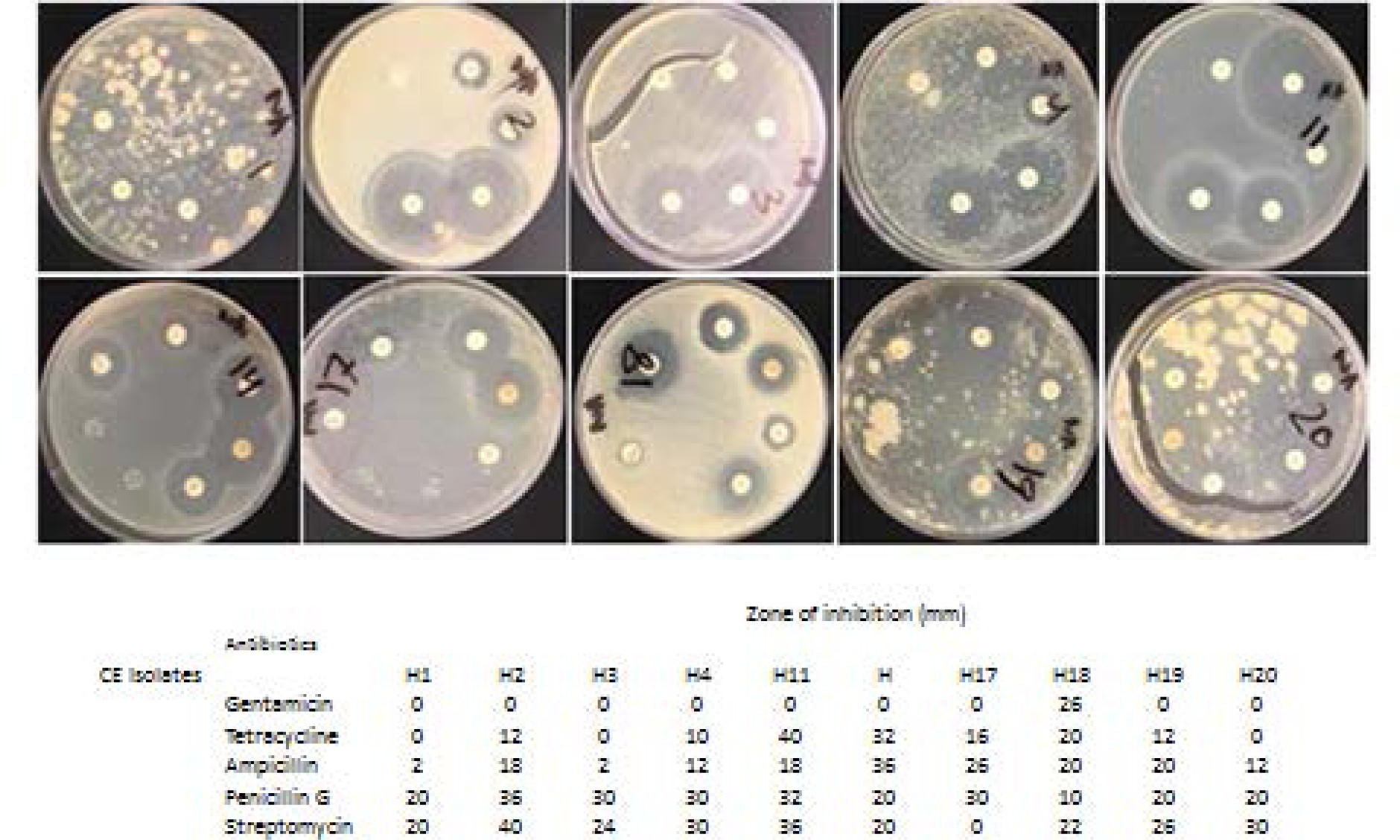
Antibiotic resistance (AR) and bacteriophage diversity in SPVs from pure culture. CE isolates revealed that isolates (H1, H2, H3, H4, and H11) did not demonstrate AR. All other isolates contained AR, however, variation in the type of AR genes were observed amongst them (Fig 6A). Interestingly, isolate CE19 contained all features required for vancomycin resistance. CE isolates were further subjected to various antibiotics that are routinely used in the clinic (eg. penicillin, ampicillin, gentamicin, streptomycin) and zone of inhibition was measured. It was found that all isolates showed some degree of susceptibility as well as resistance to different antibiotics tested (Fig 6B). In addition, Bacteriophage diversity in Spvs was also analyzed which showed all isolates tested contained at least one bacteriophage; however, variation in the type of genes was observed. Two isolates showed the presence of fungal gene (Fig 6C).

### CCR8 ligands CCL1 and CCL18 mediate adhesion, aggregation and chemotaxis of CE plaque derived T cell

It has been shown that all patients with symptomatic atherosclerotic disease show increased intraplaque lymphocytes which may be in response to pathogen stimulus. Therefore we isolated T cells (CD4+, CD8) from CE plaque and studied how these cells respond to CCL1 and CCl18. We found that T cells responded to both CCR8 ligands (CCL1 and CCL18) induction; this response was inhibited by anti CCR8 antibody (> CCR8) and inhibitor MC148 (Fig 7). CE derived T cells also formed higher number of aggregates in response to chemokine receptor inhibitors > CCR8 and MC148. This aggregation is decreased in presence of recombinant CCL1 and CCL18 suggesting that the aggregation and migration of T cells at the atherosclerotic site is mediated via CCL1/CCL18-CCR8 pathway.

**Fig 7.**
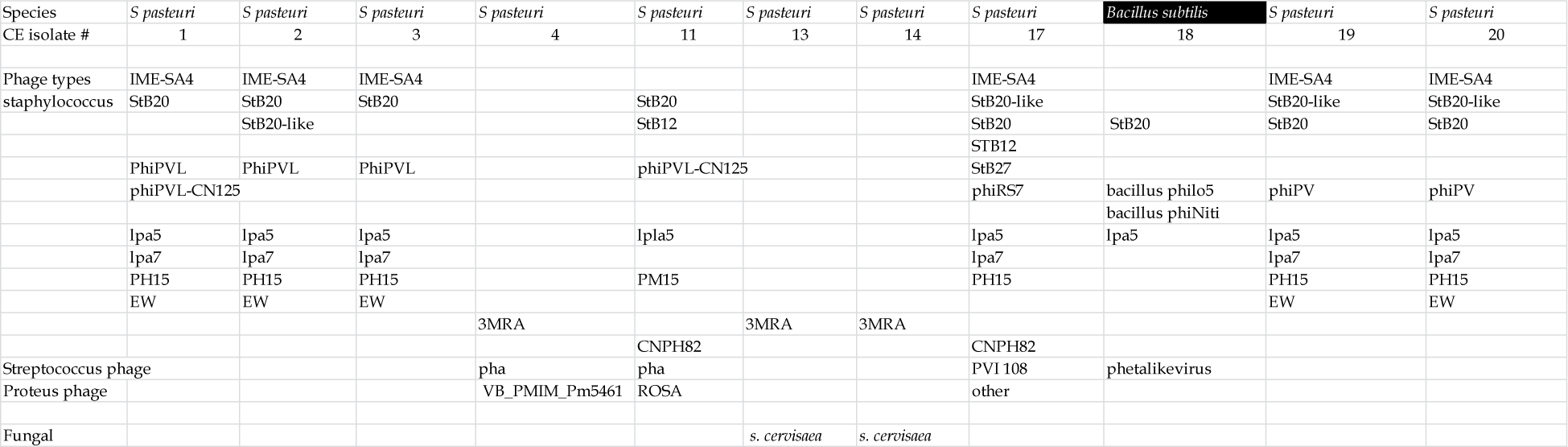
CC chemokine CCL1 and CCL18 mediate adhesion and chemotaxis of CE plaque derived T cell. T cells were isolated from CE plaque and subjected to chemotaxis to study the response to chemokine receptor CCR8 ligands, CCL1 and CCL18. IT was found that CE derived T cells (CD4+) migrated in response to both CCL1 and CCL18 and this chemotaxis was inhibited by CCR8 agonists (>CCR8 antibody and MC148). In addition CE derived T cells were found to aggregate in response to >CD3 and chemokine receptor inhibitors > CCR8 and MC148. This aggregation is decreased in presence of recombinant CCL1 and CCL18.

### *Spvs* survive phagocytosis by mouse Mφs and modulate Mφs activation, proliferation and survival

Mouse Mφs (RAW 264.7 cells) were infected with *Spvs* and it was found that *Spvs* were able to evade phagocytosis and survive within the Mφs as seen by their ability to grow on agar plate after 72 hours of culture with Mφs (Fig 8 A). *Spvs* induce Mφs activation, proliferation and survival with Spv19 having the highest capacity for inducing proliferation of Mφs in 48 hrs (Fig 8B, 8C).

**Figure 8.**
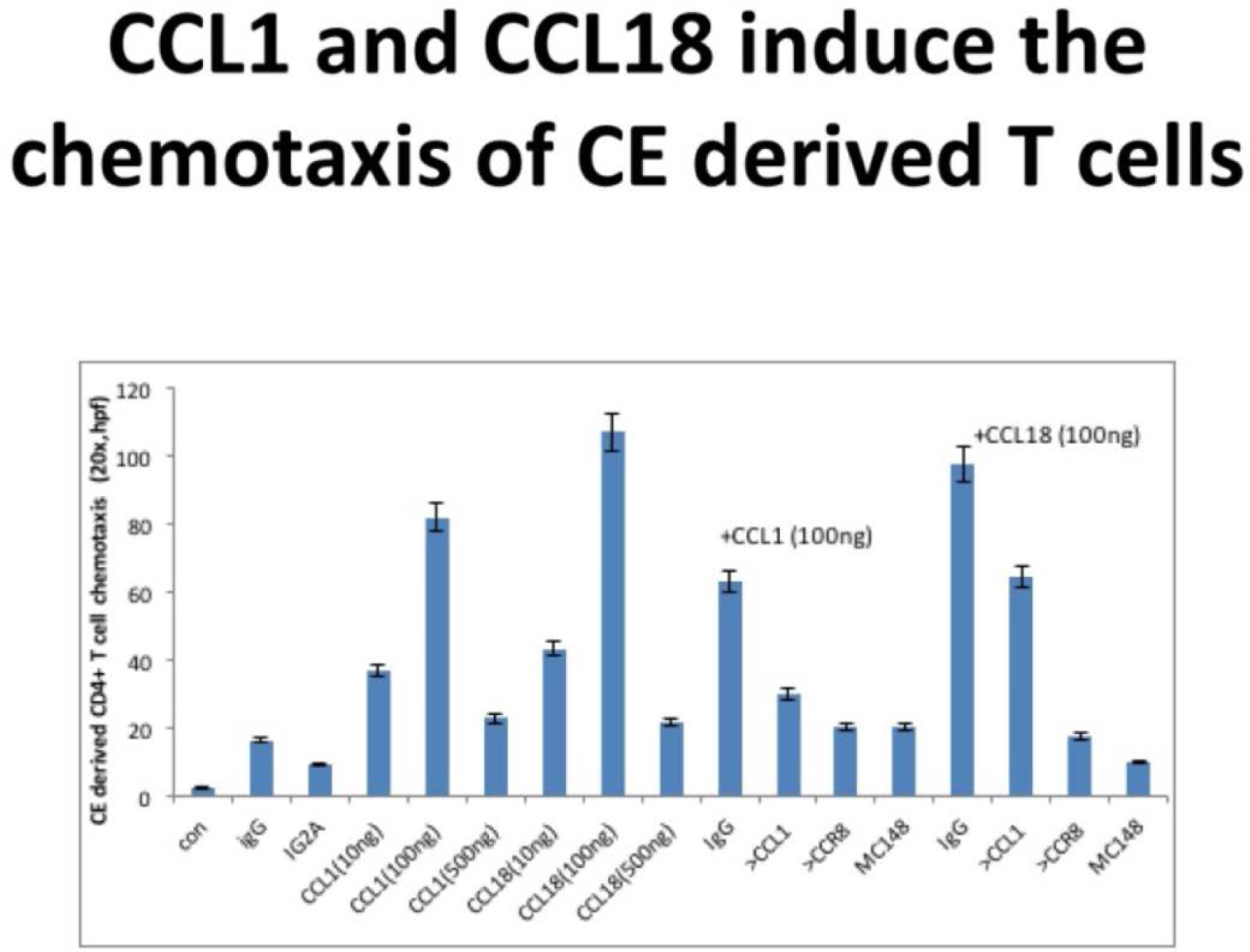
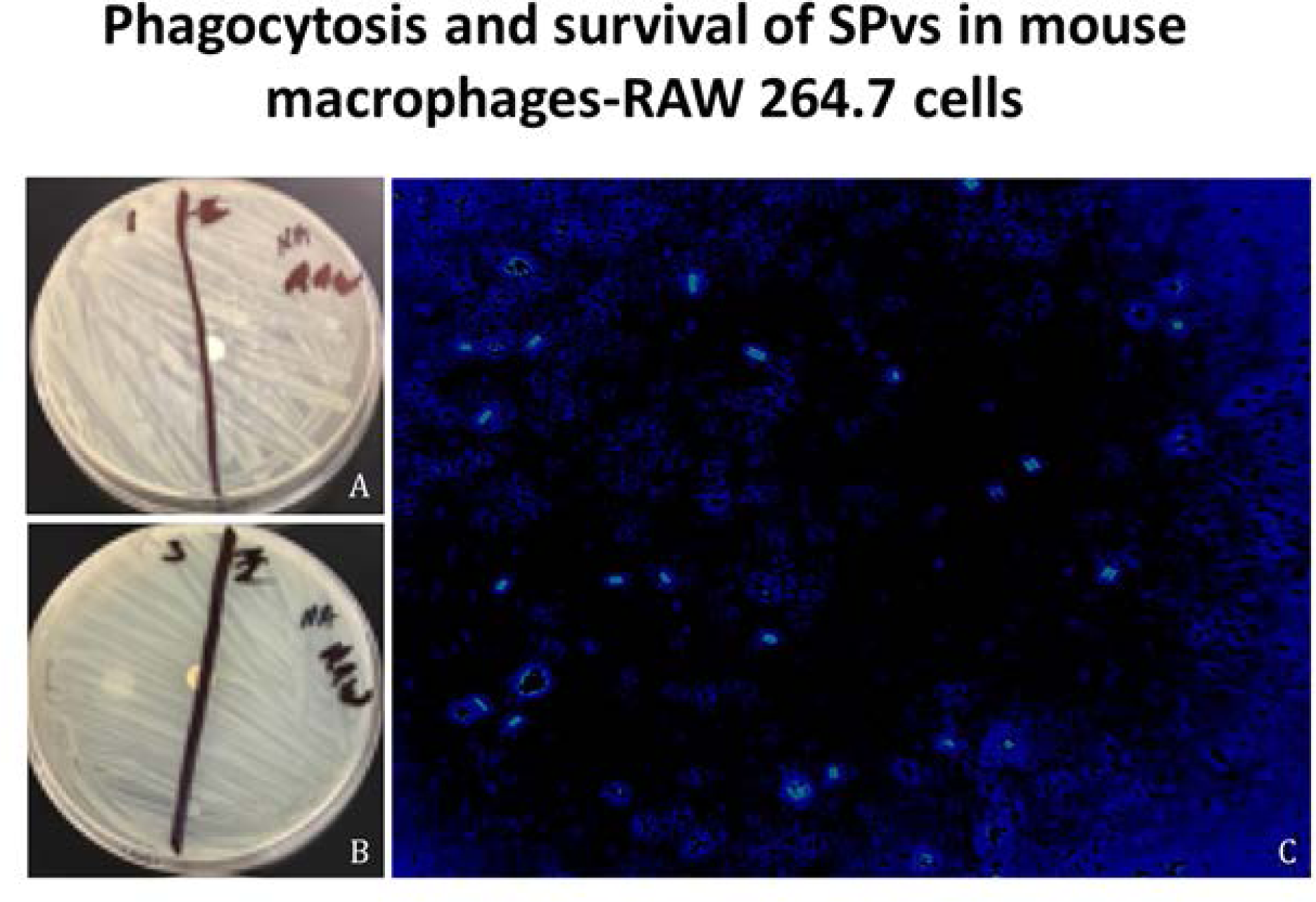
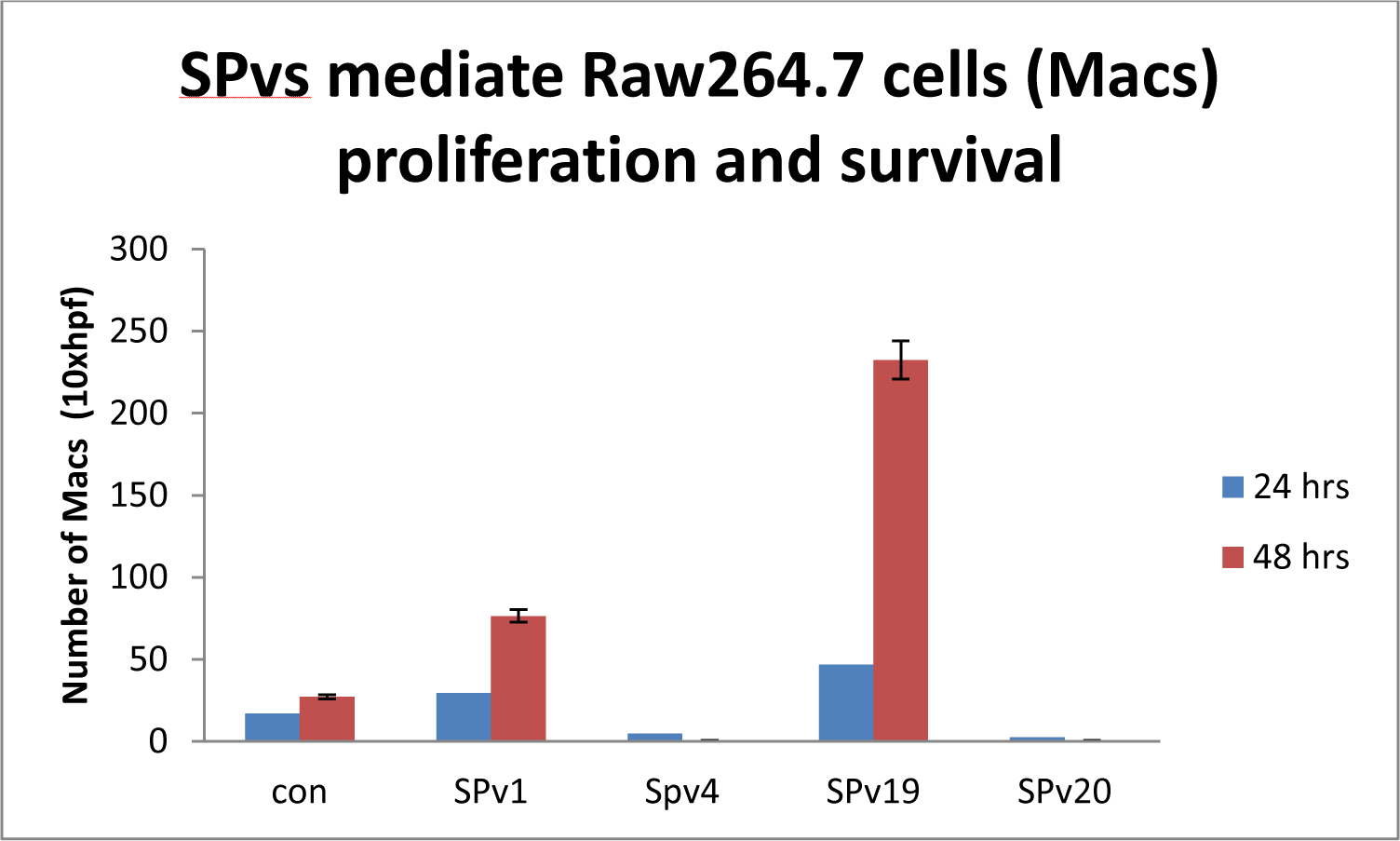
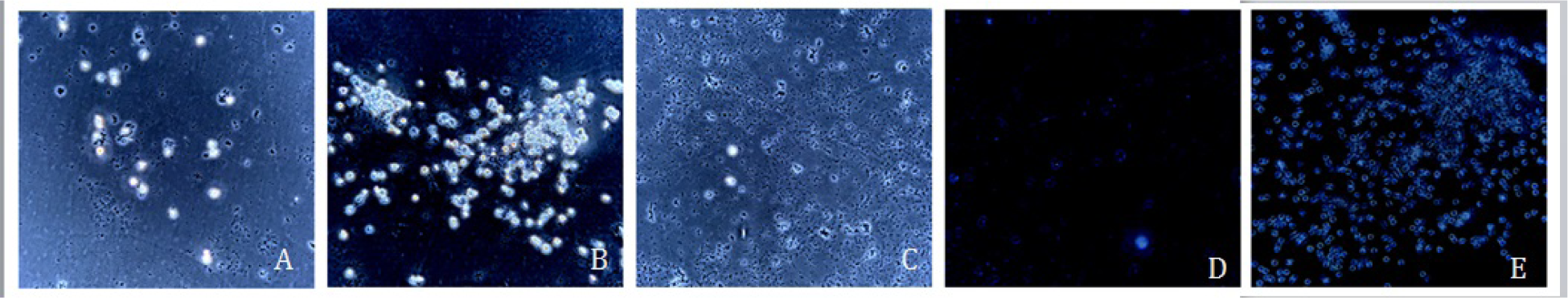
*SPvs* survive phagocytosis by mouse Mφs and modulate Mφs activation, proliferation and survival. Mouse Mφs (RAW 264.7 cells) were infected with *Spvs* and found to be able to evade phagocytosis and survive within the Mφs as seen by their ability to grow on agar plate after 72 hours of culture with Mφs [Fig 8 (a,b,c(change)]. In addition *Spvs* induce (RAW 264.7) proliferation and survival of Mφs at varying levels (Fig 8B, 8C).

### Spv19 modulation of Mφs activation is modulated via the CCL1-CCR8

Mouse Mφs were left untreated (con) or treated with CE isolates (*S. pasteuri*: *Spv1*, *Spv19* or *B. licheniformis*; *Spv18*). It was found that while *Spv19* and to a lesser degree *Spv1* activate Mφs towards which is accompanied by cell surface expression of CCR8. *Spv18* and control treatment groups do not have this effect suggesting that there is specificity to *S. pasteuri* induced via Mφ activation. *Spv19* modulation of Mφ activation is modulated via the CC chemokine and its receptor CCR8 (Fig 9)

**Figure 9.**
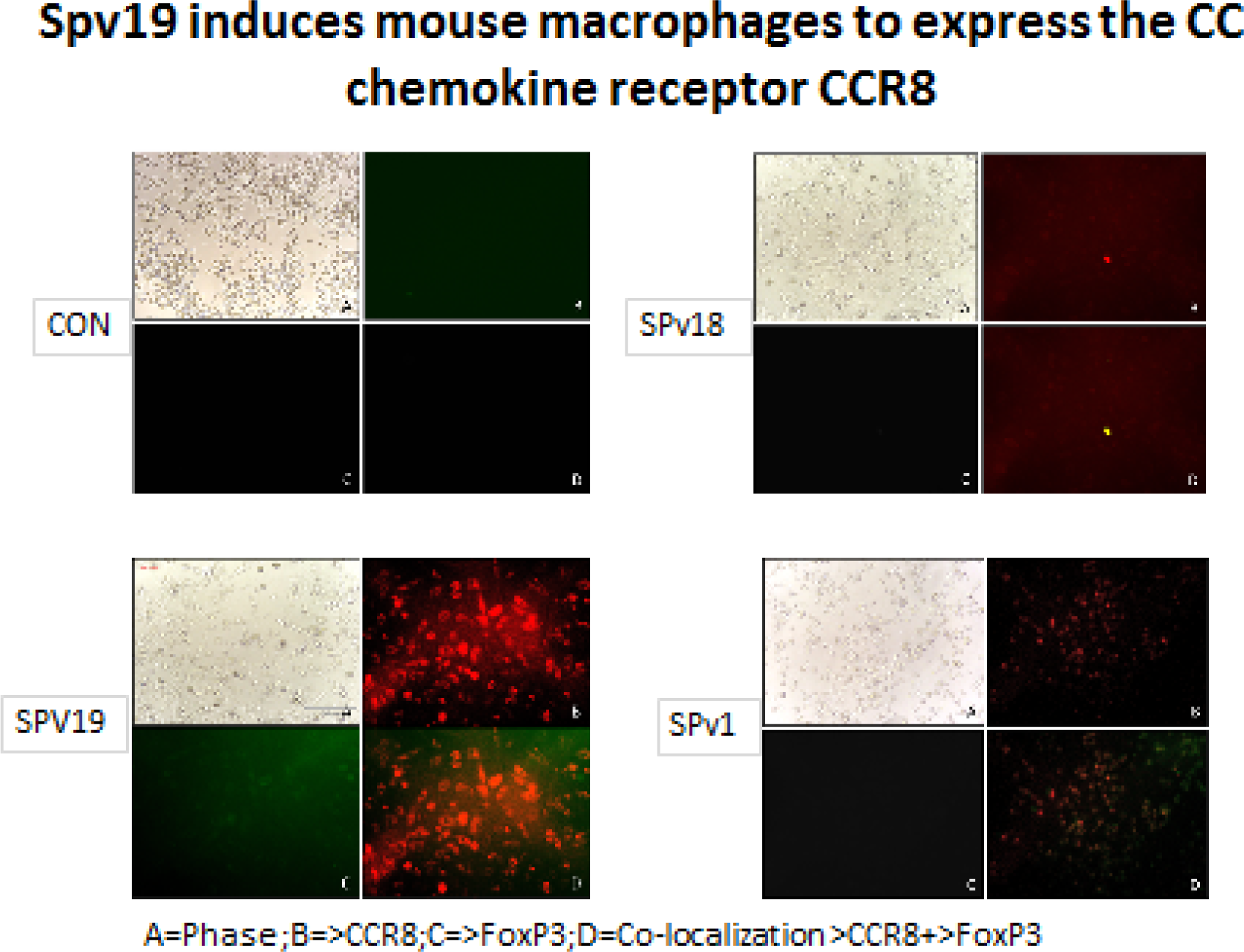
*Spv19* modulation of macrophage activation is modulated via the CC chemokine and its receptor CCR8. Mouse Mφs were left untreated (con) or treated with CE isolates (*S. pasteuri*; *Spv1*, *Spv19* or *B. licheniformis*; *Spv18*). It was found that while Spv19 and to a lesser degree *Spv1* induce conversion of Mφs towards an M1 phentotype which is accompanied by cell surface expression of CCR8. *Spv118* and control treatment groups do not have this effect suggesting that there is specificity to *S. pasteuri* induced Mφ activation.

### *Spvs* induce transmigration of Mφs across endothelial barrier

As a hallmark of atherosclerosis is endothelial denudation followed by smooth muscle proliferation and migration, we studied how *Spvs* may modulate Mφ transmigration across an endothelial barrier. Mφs were left untreated or treated with *Spv19, Spv18 and Spv17* for 48 hours; cells were washed and added to confluent human endothelial cells growing on 8 micron chemotactic chamber (Neuropore). At the end of experiment it was seen that while *Spv19* and an to a lesser extent *Spv17* modulated the trans-endothelial migration of Mφs across endothelial barrier, untreated Mφ or cells treated with *Spv18* did not demonstrate this activity (10A, 10B). In addition, the migrated cells express CCR8 showing that this was mediated via the CCL1-CCR8 pathway (Fig 10B).

**Fig 10.**
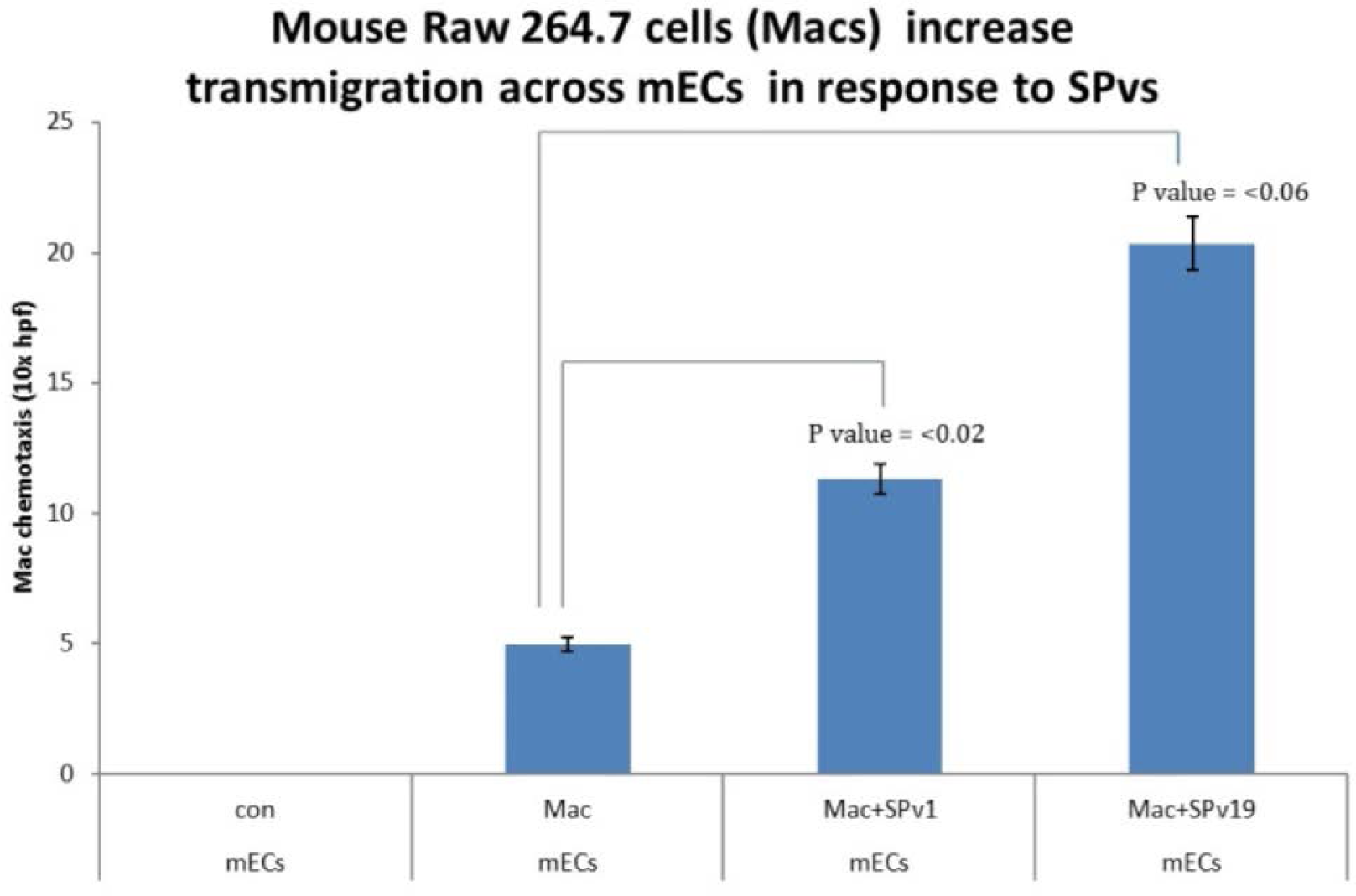
*Spv19* increases transmigration of Mφs across endothelial barrier. Mφs were untreated or treated with *Spv19* for 48 hours; cells were washed and added to confluent human endothelial cells grown on a migration chamber. It was found that while Spv19 modulated the trans-endothelial migration of Mφs across endothelial barrier untreated cells did not demonstrate this activity (10A, 10B).

### Microbial pathogenicity in atherosclerosis

A role for microbial pathogenicity in atherosclerosis is described (Box 1). Oxidation of LDL in response to iron overload and other (unknown) mediators initiates early endothelial cell injury and chemokine/cytokine signaling [e.g. Lp(a) mediated CCL1 induction], recruitment of monocyte/Mφ to the vessel wall (VW)leading to foam cell formation and initiating the and atherosclerotic process (A). Meanwhile microbial (*Spvs*) invasion of Mφs and intracellular resistance (e.g. evasion of phagocytosis) begin at distal site, chemokine receptor is activated (e.g. CCR8 in response to bacterial LPS) and microbial infiltrated Mφs are recruited to the vessel wall in response to chemokine/cytokine signals (B). The infiltrating Mφs may bring *Spvs* to the VW where bacterial colonization and biofilm formation begins and their need for iron is high (C). In order to meet their iron requirements *Spvs* scavenge heme and hemin uptake. In addition, bile hydrolysis by *Spvs* enables iron scavenging. Therefore, *Spvs* are initially tolerated at the site of injury because they reduce iron overload by scavenging of iron/heme and inhibiting the formation of oxidized LDL; thus reducing injury to ECs and subsequently to SMCs. Once the physiological source of iron has been depleted *Spvs* extract iron by and increasing production of specific siderophores. This is adaptive and the plaque remains asymptomatic. However, under stressful conditions (Host aging disease, nutrition etc.) homeostasis is disrupted. Stress response (stringent response) is initiated leading to activation of bacterial virulence and resistance. Mφ apoptosis may lead to bacterial dissemination and/or death and release of iron, heme products into the increasing oxidation of LDL and subsequent progression and destabilization of plaque and unstable symptomatic condition.

#### Box 1

##### A mechanistic paradigm for microbial pathogenicity in atherosclerosis

Microbial (*SPvs*) invasion of Mφs begin at a distal site followed by intracellular resistance and evasion of phagocytosis; activation of cytokine/chemotactic receptor (e.g. CCR8 activation in response to *Spvs* results in migration of microbial infiltrated Mφs to the site of injury (A). At the vascular wall (VW) LDL oxidation results in endothelial cell injury and recruitment of Mφ to VW (with infiltrated *Spvs*) and forms foam cells; cytokine/chemokine signaling (e.g. Lp(a) mediated CCL1 induction) results in plaque formation (B). *Spvs* scavenge iron, heme/hemin and/or increase production of siderophores to meet iron demands at the vascular wall; bile hydrolysis also increases iron scavenging as *Spvs* adapt to changing environments (C). *Spvs* **i**nitially tolerated at the site of injury due to their ability to sequester as Iron/heme reducing iron overload and limiting the formation of oxidized LDL to inhibit vascular injury and plaque remains asymptomatic. **Maladaptive:** Stress response increases bacterial virulence, resistance and Mφ apoptosis allowing bacterial dissemination/ death and release of iron, heme increasing oxidation of LDL and destabilization of plaque which is Symptomatic.

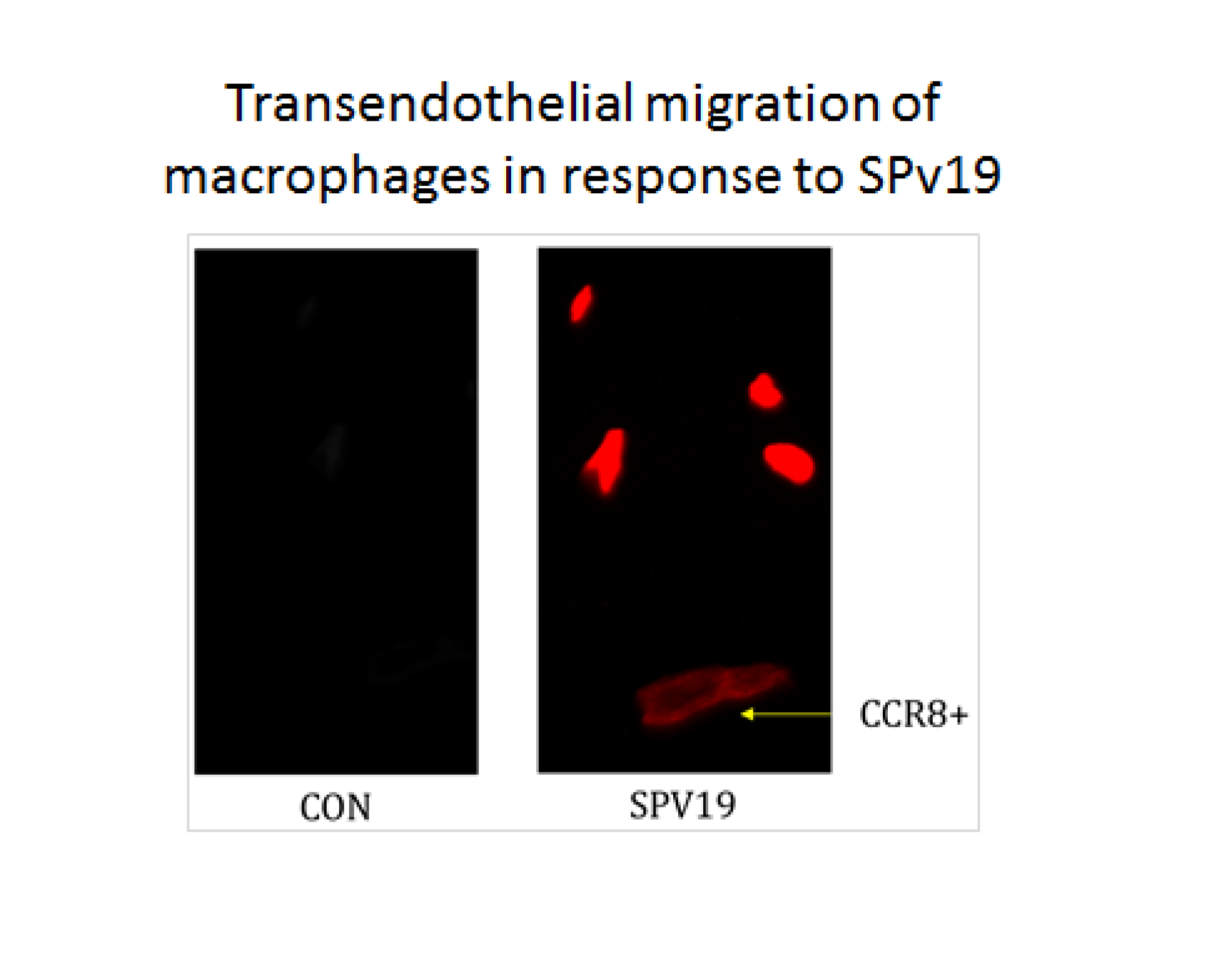

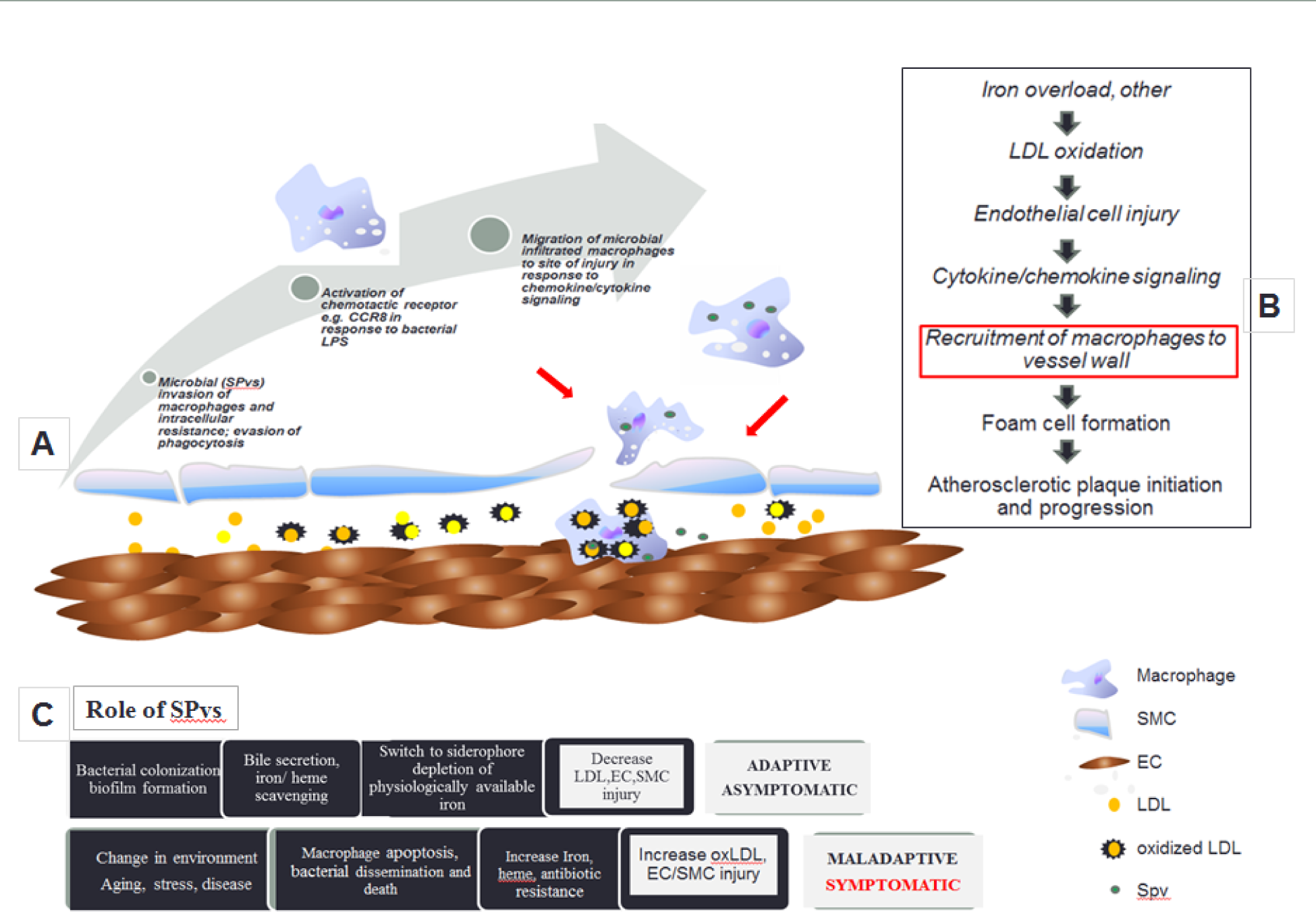

## Discussion

In this study, we have identified *Staphylococcuus pasteuri* as the predominant species in human atherosclerotic CE plaque by 16S and WGS shotgun metagenomic sequencing. We have successfully grown a series of *S. pasteuri* isolates (*Spvs*) in pure culture from CE plaque and looked for indicators of the emergence of resistances and virulence traits in this group. A repertoire of virulence has been acquired by the residents of the plaque milieu providing evidence that the CE plaque has the potential of being highly pathogenic. In addition, *Spvs* show differing levels of virulence, resistance to antibiotics and toxic compounds and are predominantly equipped for genes essential for (i) invasion and intracellular resistance and (ii) iron/heme acquisition and metabolism, which may govern Mφ invasion and subsequent homing and pathogenesis at the atherosclerotic site.

All isolates were identified as *S. pasteuri* except SPv17 which was found to be *Bacillus licheniformis*, one of the most important bacteria in industrial enzyme production of serine protease and is commonly found in the soil and on ground dwelling bird feathers and some aquatic species (23). *S. pasteuri* is a coagulase-negative, Gram positive organism which is emerging as an agent of nosocomial infections and a blood derivatives contaminant. While pathogenicity of *S pasteuri* is uncertain, it is a known contaminant in human blood of patients that causes bacteremia in humans and found to be resistant to several classes of antibiotics (methicillin/oxacillin, macrolides, lincosamides, streptogramins, tetracyclines, chloramphenicol, streptomycin, fosfomycin, as well as quaternary ammonium compounds (24). The origin of *S. pasteuri* remains unknown; however, it has been identified in routinely used substances used in human consumption such as fish (25), milk, sausage (26, 27) and drinking water (28). Its environmental presence is documented in air samples removed from the stratosphere, at an altitude of 41 km (29). In a recent study two strains of *S. pasteuri* isolates were shown to be prolific histamine formers (30). Histamine promotes the differentiation of Mφs from CD11b^+^ myeloid cells and formation of foam cells, cause damage to the arterial endothelium and modulates atherosclerosis by regulating gene expression of inflammatory modulators, an action that appears to be independent of serum cholesterol levels (31). Histamine deficiency reduces inflammation and atherosclerosis in apolipoprotein E knockout mice (32). In addition, *S pasteuri* are able to persist by physical barriers such as biofilms and/or inactivation of antibiotics secreted by other community members (33). It has been recently shown that bacteria found in the CE exist as biofilms suggesting that this may be the cause of plaque rupture turning asymptomatic plaque to become symptomatic (34).

The present study demonstrates that all *Spvs* have acquired genes for iron/heme acquisition and utilization such as those found in Gram Positives; *S aureus* reliance on heme acquisition during infection is vital to cause disease in murine models of infection [35]. While iron acquisition is necessary for bacterial survival it also plays an important role in its pathogenesis, however, it is the only nutrient not freely available to bacteria in the human host [<u>36</u>]. To circumvent this, pathogens have evolved elaborate strategies to acquire iron via regulating gene expression to produce accessory toxins, siderophores and/or hemolysins etc. Oxidation of low density lipoprotein (LDL) by physiological substances such as iron/heme is known to play a key role in the initiation and progression of atherosclerosis. This is supported by the finding that LDL oxidized by heme is extremely cytotoxic to cultured aortic endothelial cells that may also have further implications for similar oxidative reactions between heme and unsaturated fatty acids which may occur consequent to hemorrhagic injury (37). Thus, iron/heme sequestration from the plaque milieu by *Spvs* may, in part, be responsible for circumventing endothelial damage and limiting detrimental plaque progression. In addition, we found that *Spvs* have acquired genes for bile hydrolysis which is known to increase iron scavenging. Bile salts increase mRNA levels for genes associated with iron scavenging and metabolism which is counteracted by the inhibitory effect of the iron chelating agent 2,2’-dipyridyl on growth of enteric pathogen, *Escherichia coli* O157:H7 suggesting that this organism may use bile as an environmental signal to adapt to changing conditions associated with the small intestine, including adaptation to an iron-scarce environment (38). Moreover, bile exposure induced significant changes in mRNA levels of genes related to virulence potential, including a reduction of mRNA for genes making up the locus of enterocyte effacement (LEE) pathogenicity island. We show that *Spv1* and *Spv19* have acquired a pathogenicity island found in *E. coli* in the LEE (EPEC) known for forming attaching and effacing lesions (39). Whether *Spvs* virulence and release of iron can trigger formation of unstable plaques remains an interesting possibility.

Among all the CE isolates *Spv19* was found to contain the highest number of genes for cell wall synthesis to support its requirement to maintain its growth, development and reproduction. Interestingly, *Spv13* with low number of genes for cell wall synthesis contained the highest number of genes for secondary metabolites compared to all isolates. The results suggest that key metabolic activities that are essential for pathogenesis might be dispensable during persistent infections and sub-clinical infection may be a key driver governing the relative representation of *Spvs* in the plaque due to a stringent response (SR) adaptation evolved in response to negative selection pressures. The SR is a bacterial response to a multitude of different environmental stress conditions which is characterized by the synthesis of the messenger molecules (p)ppGpp which also play a key role for pathogens to switch between specific phenotypic states within the host (42). While stringent response is unknown for *S. pasteuri*, stringent response in another staphylococcus strain *S. aureus* is well established. Notably, Staphylococci are known to express a great variability and flexibility in metabolic expression to adapt to different environmental conditions and survive (43). For example, killing of phagocytes as well as survival within these cells has been proposed as a major mechanism for the success of *S. aureus* to spread within the body (44); thus activating the virulent phenotype by the stringent response. This suggests that this species may utilize its virulence for multiple tasks depending on the stage/state of development of the atherosclerotic plaque. Clonal hematopoiesis of intermediate potential is a known risk factor in tumor pathology and has recently been shown to be higher in individuals with atherosclerosis (45, 46). Variation in host stem cells (due to aging, disease) may influence differential immune response to pathogenic bacteria at the site of injury which may influence the mediation of symptomatic vs. asymptomatic atherosclerosis. For example, age induced somatic mutation may enhance the pathogen’s ability for invasion and colonization.

As Mφs are known to contribute to the initiation, development and progression of atherogenesis, we infected mouse Mφs with *Spvs* to study its role in the modulation of these immune cells. We have previously shown that the CC chemokine and its receptor CCR8 are found on human atherosclerotic plaque, modulate monocyte, VSMC and EC migration and proliferation in response to Lipoprotein (a) [Lpa(a)] (40,41). In the present study we show that *Spvs* differ in their ability to modulate mouse Mφ activation, proliferation, migration, differentiation and survival of mouse Mφs which is mediated by CCL1/CCR8. Spvs from different patients varied in eliciting host immune via CCR8 activation on Mφs suggesting that *Spvs* may also contribute to Mφ modulation and consequently to pathogenesis. For example, CE plaque derived T cell adhesion, aggregation and migration is mediated via the CCL1-CCR8 pathway. In addition, we show that *SPv1* and *SPv19* survive phagocytosis by mouse Mφs and modulate trans-endothelial cell migration of these cells. None of the *Spvs* had any motility or chemotactic genes yet were found at the site of the plaque suggesting that Mφ invasion is the primary mode of transport to the site of injury. Interestingly, *B. licheniformis* was the only organism found in our study to have the ability of independent motility which is shown by the abundant supply of flagella genes.

In conclusion, we show that *S. pasteuri* isolated from the CE plaque contain genes for virulence with the ability to escape phagocytosis, respond to pathogenic signals and transmigrate across an endothelial barrier. This process is mediated via CCL1-CCR8 pathway which we have previously demonstrated as a modulator of the pathophysiology of atherosclerosis (41,47). To the best of our knowledge, this is the first study that provides a detailed genomic analysis of *S. pasteuri* isolates found in human atherosclerotic tissue and offers a novel paradigm for microbial pathogenicity in atherosclerosis (Box1). Although *S. pasteuri* has previously been found in human blood, this is the first demonstration of *S. pasteuri* and its variants (*Spvs*) in the atherosclerotic plaque. Virulence in *Spvs* demonstrated in this study is most likely facilitated by concerted activities of mobile genetic elements which facilitate horizontal genetic exchange and therefore promote the acquisition and spread of virulence. They are bound to be other pathogens that may play pivotal roles in the development and/or progression of this disease. The generalists’ and specialists in the colonizing microbiome at the site of injury, the circumstance of their activation/SD and the host response must be understood. What we may find is a common pathogen, an etiological agent of atherosclerosis; or, what is more likely, a repertoire of virulent genes acquired over a lifetime of an individual; a unique evolutionary trajectory for each individual. While limited by the analysis of a single isolate at each time point, this study adds to the available information regarding the characteristics of atherosclerosis and highlights the significance of carotid vessels as a reservoir for *S. pasteuri* and the ability of this pathogen to persist for long periods which is most likely associated with its conserved virulence. Low cost of sequencing and advances in NGS technology should allow clinical studies, across multiple ethnicity and geographical locations which is warranted for a study of this magnitude if we are to make any attempt at defining the atherosclerotic microbiome. In future, novel tools need to be developed in order to characterize the host-microbe relationships in the atherosclerotic niche which may offer novel insights into the understanding of this disease. Thus, coupling microbial virulence factors to the atherosclerotic process may explain the persistent infection in atherosclerosis.

## Supplemental data

All supplemental data is provided in separate file which included Figures S1 S2, S3, S4, S5, S6,S7 and S8.

