## Supplemental data for "*Staphylococcus pasteuri* isolates (*Spvs*) from human atherosclerotic plaques mediate virulence, intracellular resistance and transendothelial invasion of macrophages: A mechanistic paradigm for microbial pathogenicity in atherosclerosis"

**Supplemental figures**

Fig S1

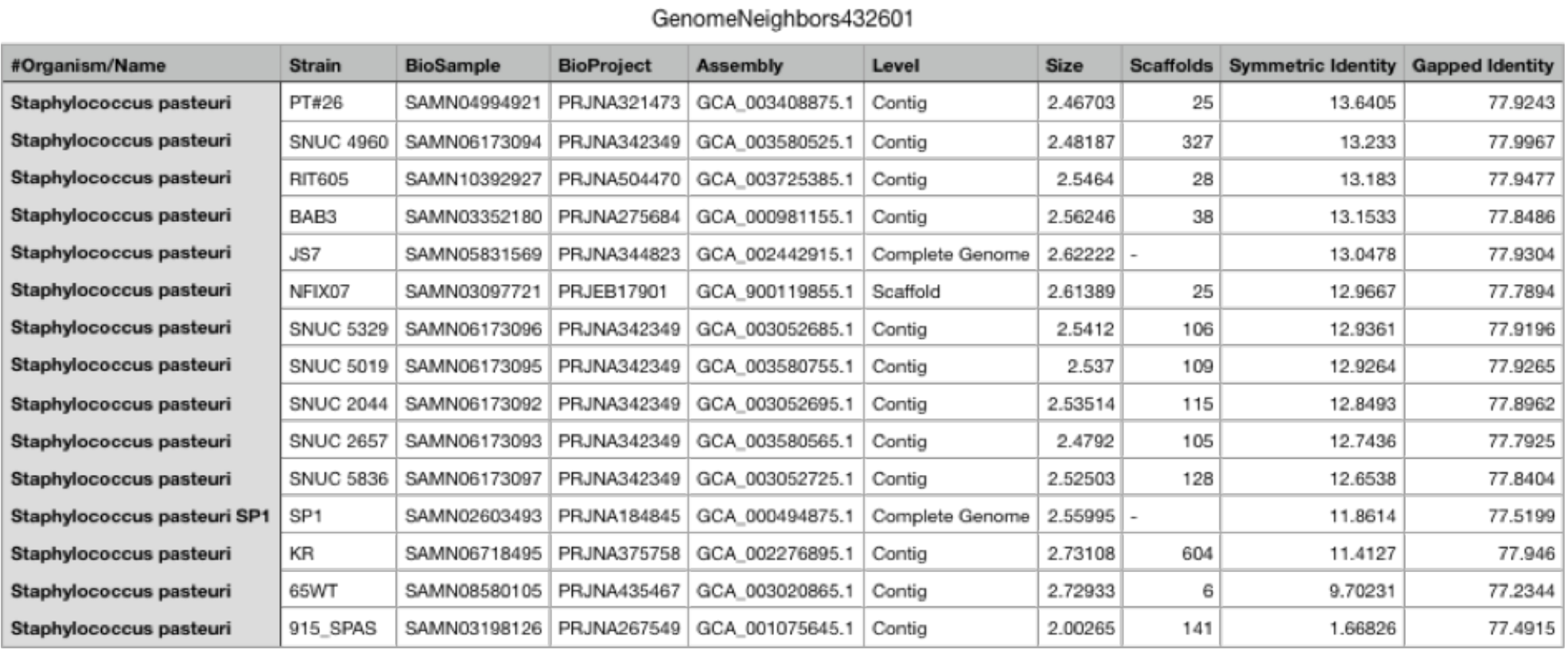

Fig S2

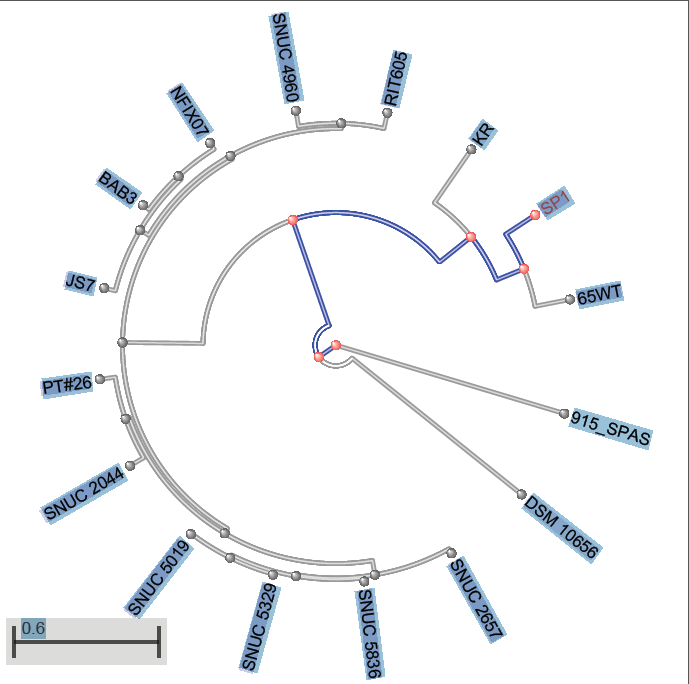

Fig S3

(*Spv18*)

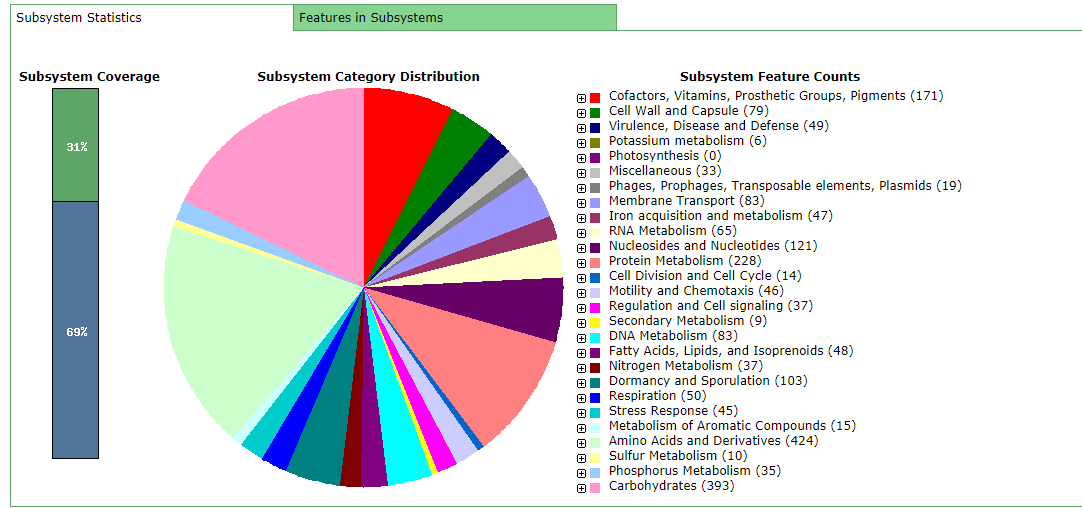

Fig S4

(*Spv17*)

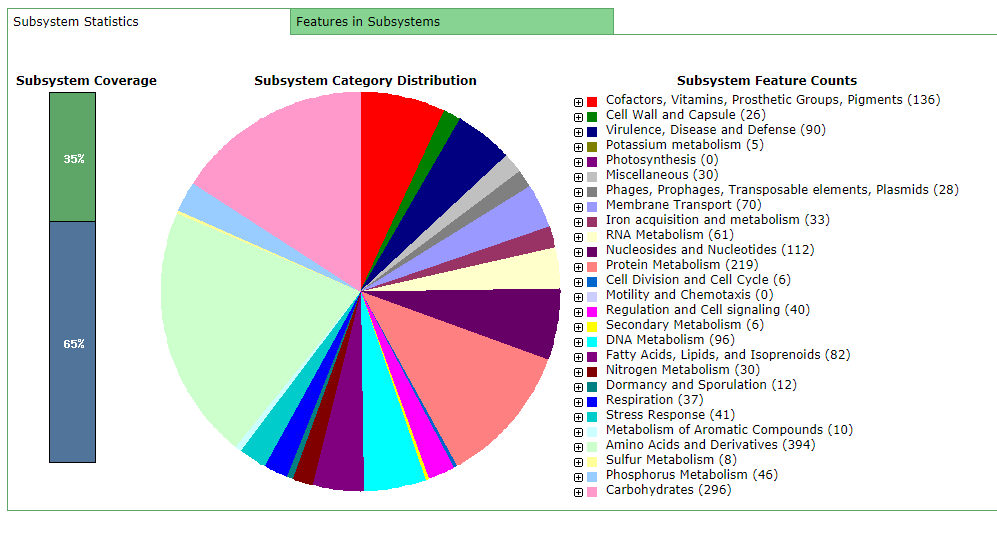

Fig S5

(*Spv14*)

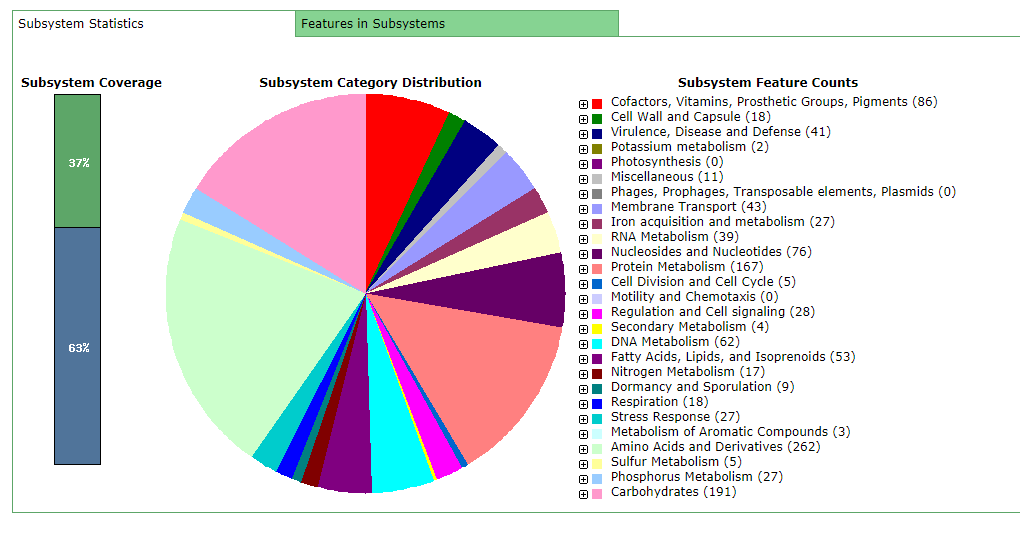

Fig S6

(*Spv13*)

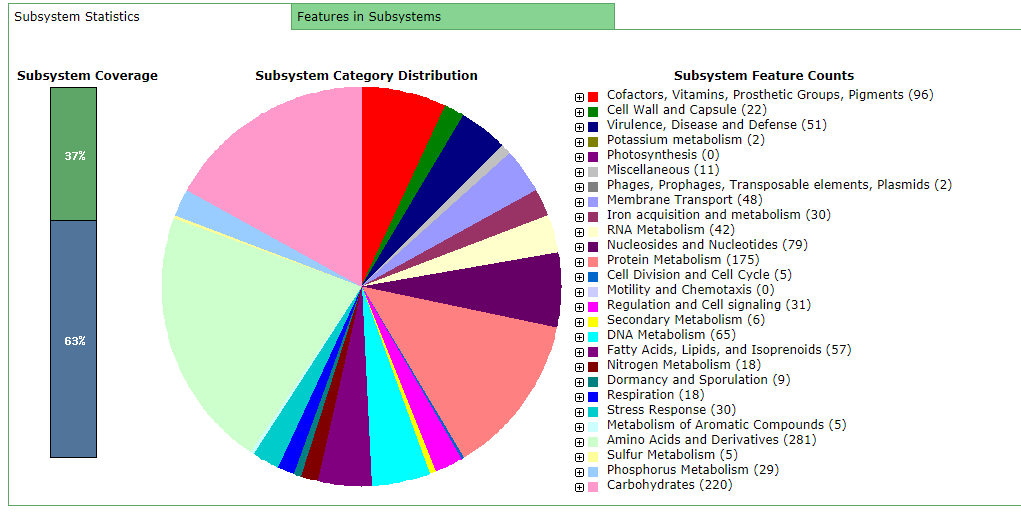

;;

Fig S7

(*Spv11*)

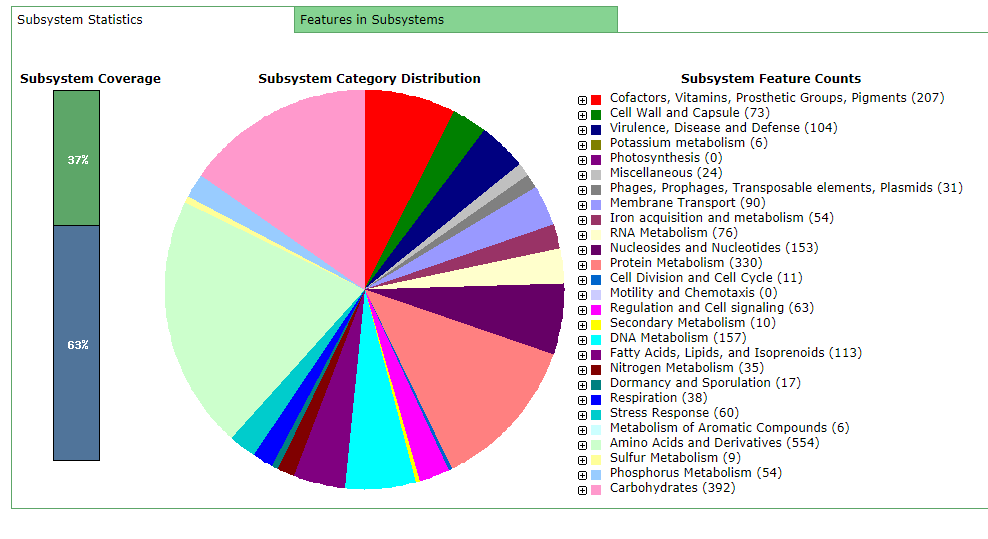

Fig S8

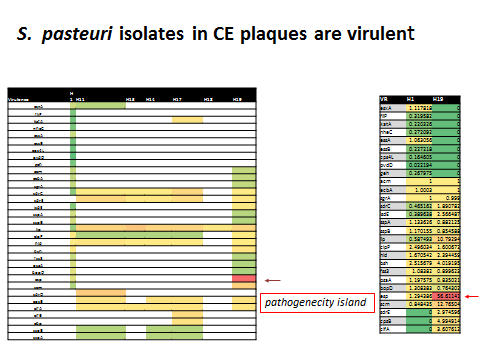

**Legend - Supplemental figures**

S1 Genome

S2 Tree

S3 H18 subsystems

S4 H17 subsystems

S5 H14 subsystems

S6 H13 subsystems

S7 H11 subsystems

S8 Virulence in different *Spv* isolates

S8	*S. pasteuri* isolates in CE plaques are virulent with abundance of E. coli derived Pathogenic

** Supplementary figure S8 (accompanied with pathogenicity) island*

**All data will be deposited to GenBank on submission of manuscript for publication**
